# Generation of Region-specific Airway Basal Stem Cells from Human Pluripotent Stem Cells via Regulation of NOGGIN-BMP Axis

**DOI:** 10.1101/2025.10.01.657190

**Authors:** Shingo Suzuki, Nicole V. Acosta Sandoval, Shnutez Doddipalli, Dalia M. Hassan, Cristina Barillà, Andras Rab, Mo-Fan Huang, John M. Avila, Barbara Tabak, Junjie Lu, Peggy P. Hsu, Tristan Frum, Bryan C. Derebery, Maria Gacha-Garay, Andres D. Chamorro-Parejo, Samantha L. Winkler, Sadhana Srinagasai Ponnaluri, Nicolas Forcioli-Conti, Martha Rea-Moreno, Jinho Kim, Ciaran M. Lee, Naoki Nakayama, Wenbo Li, Ya-Wen Chen, Kenneth S. Ramos, Jason R. Spence, John E. Mahoney, Dung-Fang Lee, Eric J. Sorscher, Jichao Chen, Brian R. Davis, Sarah X. Huang

## Abstract

Basal cells (BCs) are the primary stem cell population of adult human airways and a key target for hPSC-based models of airway development, disease and regenerative medicine. Recent studies have revealed substantial regional differences between human proximal and distal airway cell types, including BCs. Here, we show that the NOGGIN-BMP signaling axis governs proximal-distal patterning of hPSC-derived lung progenitors leading to the generation of region-specific induced BCs (iBCs). Continuous BMP inhibition through NOGGIN, coupled with tapered WNT activation, generates proximal iBCs that molecularly and functionally resemble human proximal airway BCs and differentiate *in vitro* and *in vivo* into the full repertoire of specialized proximal airway cell types, including ionocytes and pulmonary neuroendocrine cells. In sharp contrast, BMP activation with tapered WNT generates distal airway-like cells and distal BCs with limited ionocyte differentiation potential. Progeny of proximal iBCs derived from G551D *CFTR* mutant hiPSCs also recapitulate a cystic fibrosis-associated ionocyte phenotype.

## INTRODUCTION

Airway basal cells (BCs) are small cuboidal epithelial cells located adjacent to the basal lamina of the surface airway. Adult airway BCs express canonical markers including Tumor protein 63 (TP63), cytoskeletal protein keratin 5 (KRT5), and nerve growth factor receptor (NGFR). In both mouse and human systems, BCs are established adult airway stem cells with self-renewal and multilineage differentiation capacity, making them promising candidates for cell-based therapies and models of airway diseases such as cystic fibrosis (CF).

While BCs were once considered homogeneous, recent studies have revealed marked proximal-distal heterogeneity among airway epithelial cell types, including BCs, in abundance, molecular phenotype, and function (Montoro et al. 2018; Plasschaert et al. 2018; Okuda et al. 2020; Travaglini et al. 2020; Kadur Lakshminarasimha Murthy et al. 2022; He et al. 2022). In terms of abundance, BCs in mice are largely restricted to the trachea, whereas in humans they extend from the cartilaginous trachea and proximal conducting airways into non-cartilaginous intermediate and distal airways. Secretory and multiciliated cells are distributed throughout the conducting airways in both species, whereas rare cell types such as ionocytes and tuft (brush) cells are enriched in the trachea and proximal cartilaginous airways across species (Montoro et al. 2018; Okuda et al. 2020; Plasschaert et al. 2018; Travaglini et al. 2020; Yuan et al. 2023). Phenotypically, basal and secretory cells exhibit region-specific features along the proximal–intermediate–distal axis of human airways (Kadur Lakshminarasimha Murthy et al. 2022). Functionally, human proximal and distal airway BCs retain intrinsic region-specific properties *ex vivo*, differentiating into progeny corresponding to their respective anatomical regions (Kadur Lakshminarasimha Murthy et al. 2022; Zhou et al. 2022). Together, these findings underscore the need to define the region-specific roles of BCs, as the major airway stem cell population, in lung homeostasis, repair, and disease.

Because mouse BCs are largely confined to the trachea and scarce in lower airways, and ferret models are costly and not widely available, studies of regional BC heterogeneity currently rely largely on primary human BCs (Suprynowicz et al. 2012; Mou et al. 2016), which have limited accessibility and expansion potential. Thus, a renewable source of region-specific human BCs is critically needed. Human pluripotent stem cells (hPSCs), including patient-derived human induced pluripotent stem cells (hiPSCs) and human embryonic stem cells (hESCs), can potentially be differentiated into region-specific BCs, providing a scalable renewable cell source.

Differentiation of hPSCs into airway BCs begins with the generation of lung progenitors expressing NK2 homeobox 1 (NKX2-1), the defining transcription factor of the lung epithelial lineage. Multiple groups, including ours, have developed methods to derive NKX2-1^+^ lung progenitors from hPSCs (Hawkins et al. 2017; Huang et al. 2014; McCauley et al. 2017; Rankin et al. 2016; Konishi et al. 2016; Yamamoto et al. 2017; Dye et al. 2015; Mou et al. 2012; Firth et al. 2014; Wong et al. 2012), which can further differentiate into both airway and alveolar lineages (Mou et al. 2012; Huang et al. 2014; Hawkins et al. 2017; Dye et al. 2015; Chen et al. 2017). However, prior studies have neither deliberately modulated proximal-distal patterning during subsequent airway differentiation nor systematically characterized the regional identity of the resulting lineages.

Recent efforts have aimed to enrich airway epithelial cell types and BCs during hiPSC differentiation. Guided by mouse studies (Hou et al. 2019; Hashimoto et al. 2012; Mucenski et al. 2003; Shu et al. 2005), WNT downregulation has been used to direct hiPSC-derived lung progenitors toward airway specification (McCauley et al. 2017). Using this strategy, putative basal progenitors expressing NKX2-1, TP63, and KRT5 can be purified and matured into induced BCs (iBCs) in medium optimized for adult BC expansion (Hawkins et al. 2021). These iBCs successfully differentiate into secretory and multiciliated cells *in vitro* and *in vivo*, but rare cell output, particularly ionocytes and pulmonary neuroendocrine cells (PNECs) differentiation, has been limited (Hawkins et al. 2021) and was mainly reported in a subsequent study after repeated passaging (Wang et al. 2023). Another study adapted the WNT-downregulation method to generate mixed airway lineages from hiPSCs without basal-cell purification and observed sporadic ionocytes in subsequent *in vitro* differentiation cultures (Vila-Gonzalez et al. 2024). Because these prior studies did not explicitly assess regional airway identity, it remains unclear whether the resulting iBCs correspond to proximal, intermediate, or distal airway BCs, or a mixture of these identities. Together with the sporadic or variable ionocyte output and the secretory phenotypes reported in prior iBC studies (Hawkins et al. 2021; Wang et al. 2023), these findings suggest that previously generated iBCs predominantly adopt an intermediate-airway identity. Thus, the directed differentiation of hPSCs toward region-specific iBCs, particularly proximal iBCs, remains an unmet need.

In this study, we demonstrate successful differentiation of hPSCs into region-specific iBCs that closely recapitulate the molecular and functional phenotypes of human adult proximal, intermediate, and distal airway BCs. We achieve this by recapitulating key signaling events involved in proximal-distal patterning, airway basal specification, and lung fate maintenance. Specifically, we identify the NOGGIN–bone morphogenetic protein (BMP) signaling axis as a critical regulator of proximal-distal patterning during airway basal differentiation. This finding aligns with and extends prior mouse and human fetal lung studies, distinguishing the role of BMP signaling from other pathways implicated in proximal-distal patterning, including WNT, fibroblast growth factor (FGF), transforming growth factor beta (TGF-β), NOTCH, retinoic acid (RA), and epidermal growth factor (EGF) signaling (Hashimoto et al. 2012; Mucenski et al. 2003; Shu et al. 2005; Bellusci et al. 1997; Hyatt, Shangguan, and Shannon 2004; Bellusci et al. 1996; Lu et al. 2001; Weaver et al. 1999; Miller et al. 2020). In parallel, we define a critical role for WNT signaling in maintaining lung fate during basal cell differentiation. Prior studies indicate that WNT downregulation is required for airway basal specification (Hou et al. 2019; Hashimoto et al. 2012; Mucenski et al. 2003; Shu et al. 2005; McCauley et al. 2017; Hawkins et al. 2021), but not a major determinant of proximal-distal regional patterning (McCauley et al. 2017; Hawkins et al. 2021). However, premature WNT withdrawal can lead to unintended loss of NKX2-1 lung fate (McCauley et al. 2017; Hawkins et al. 2021), consistent with our previous finding that tonic WNT signaling maintains lung fate after initial NKX2-1^+^ lung progenitor specification in both humans and mice (Ostrin et al. 2018). In this study, our approach tapers WNT signaling to balance the WNT reduction required for basal specification with the activation needed to maintain lung fate, thereby recapitulating the temporal shift toward WNT-independent NKX2-1 maintenance observed *in vivo* (Ostrin et al. 2018; Gerner-Mauro, Akiyama, and Chen 2020).

Here, we show that after lung specification, continuous NOGGIN-mediated BMP inhibition combined with tapered WNT signaling directs progenitors toward a proximal basal fate, whereas BMP activation with tapered WNT signaling promotes distal iBC differentiation. Proximal iBCs robustly differentiate into ionocytes and secretory cells with proximal phenotypes *in vitro* and *in vivo*, while distal iBCs show minimal ionocyte potential and secretory phenotypes characteristic of small airways. Both proximal and distal iBCs exhibit long-term self-renewal in defined serum-free medium, and both iBCs and their progenitors can be cryopreserved at multiple stages while retaining their regional identity. These region-specific iBCs provide a platform for studies of airway development, rare cell biology, and disease, including modeling the ionocyte phenotype associated with the G551D *CFTR* mutation in CF.

## RESULTS

### A four-stage differentiation strategy that recapitulates *in vivo* development

We derived region-specific iBCs using a four-stage differentiation strategy (Figure 1A) that recapitulates sequential milestones of *in vivo* lung development. Stage 1 generated lung progenitors resembling early primordial NKX2-1^+^ lung buds, which express sex-determining region Y-box 9 (Sox9) in mice and are SOX2^low^SOX9^+^ in humans. In Stage 2, we generated proximal airway progenitors and intermediate/distal multipotent progenitors, respectively. At this stage, the basal transcription factor TP63 is not yet activated because lung fate maintenance requires continued high-level WNT activation, as in Stage 1 lung progenitor specification (Ostrin et al. 2018; Gerner-Mauro, Akiyama, and Chen 2020), whereas basal specification requires reduced WNT signaling (Hou et al. 2019; Hashimoto et al. 2012; Mucenski et al. 2003; Shu et al. 2005). In Stage 3, these airway/multipotent progenitors were directed toward putative proximal, intermediate, and distal airway basal progenitors expressing TP63. These progenitors were subsequently maintained and matured during Stage 4 to express canonical basal markers KRT5 and NGFR. NGFR, a mature basal cell surface marker expressed in postnatal mouse and human airway epithelium (Rock et al. 2009), was used at this stage to quantify basal maturation efficiency across conditions.

**Fig 1.**
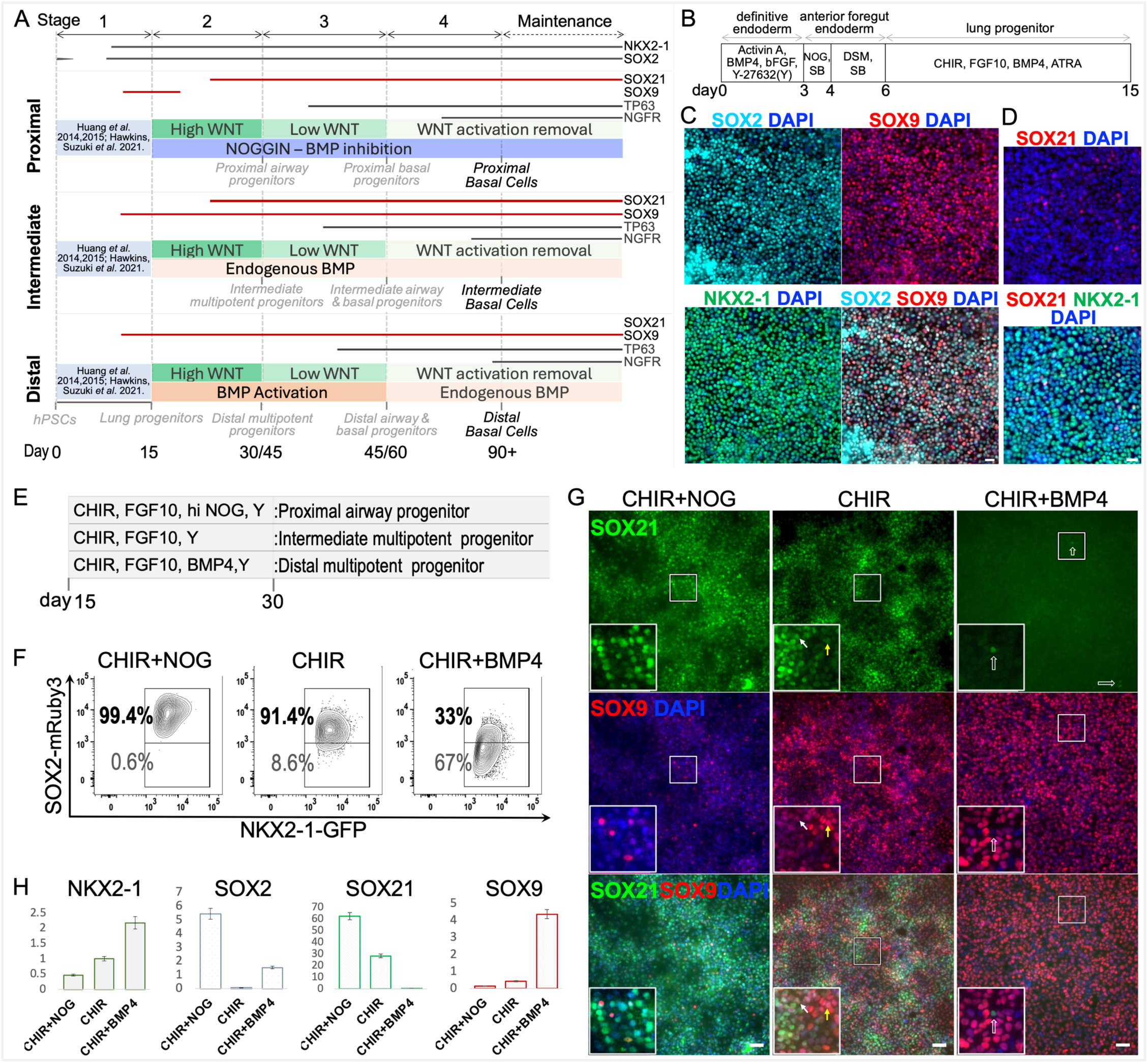
NOGGIN-BMP signaling systematically regulates SOX2, SOX21, and SOX9 expression in hPSC-derived lung progenitors. (**A**) Schematic illustration of the overall differentiation strategy. Not drawn to scale. (**B**) Schematic of the culture protocol for days 0–15 (stage 1). Y, Y-27632; NOG, NOGGIN; DSM, Dorsomorphin dihydrochloride; SB, SB431542; CHIR, CHIR99021, a pharmacological agonist of WNT signaling; ATRA, all-trans retinoic acid. (**C–D**) Expression of SOX2/SOX9/NKX2-1 (C) and SOX21/NKX2-1 (D) in cells cultured according to the protocol shown in B. Scale bar, 20μm. n = 3 for each hPSC line, representative images from hiPSC line BU3NGPT. (**E**) Schematic of the culture protocol for days 15–30 (stage 2). hi NOG = high NOGGIN. **(F)** SOX2^mRUBY3^ reporter protein profiles in NKX2-1^GFP+^ R2 NGSmR cells cultured according to E. CHIR + NOG: Proximal; CHIR: Intermediate; CHIR + BMP4: Distal. (**G**) SOX21 and SOX9 protein expression in NKX2-1^GFP+^ cells by IF staining from indicated conditions cultured according to E. Cells were extracted from 3D culture, NKX2-1^GFP+^ cells were sorted and briefly attached on 2D for 24–48 hours before staining. White and yellow arrows mark representative SOX21⁺SOX9^low^ and SOX21^low^SOX9⁺ cells, respectively, under the intermediate condition; open white arrows indicate sporadic SOX21⁺ cells under the distal condition. Scale bar, 50μm. *n* = 3 for each hPSC line, representative images from hiPSC line BU3 NGPT. (**H**) Verification of *NKX2-1, SOX2, SOX21* and *SOX9* mRNA expression by qPCR under indicated conditions of NKX2-1^GFP+^ BU3 NGPT cells cultured according to E. The expression levels were shown normalized relative to an alveolar progenitor-enriched condition as reported in our previous publication (Huang *et al*. 2014), *n* = 3, a representative experiment is shown.

To assess proximal-distal regional identity during Stages 2–4, we established a developmental patterning roadmap based on published evidence and our analysis of human fetal lung (HFL). In mice, branching morphogenesis segregates Sox2^+^ airway stalks from Sox9^+^ distal tips (Hashimoto et al. 2012; Liu and Hogan 2002). In humans, early distal tips are SOX2^low^SOX9^+^ and gradually turn off SOX2 by gestation week (GW) 21, while airway stalks become SOX2^+^ (Xu et al. 2016; Nikolic et al. 2017). However, because all airway progenitors express SOX2, this information alone was insufficient to guide regional airway patterning during our differentiation; therefore, we incorporated SOX21 to further resolve proximal airway identity. In mice, Sox21 predominantly marks the most proximal Sox2^+^ airways – the trachea and main bronchi (zone 1), while zone 2 cells in the intrapulmonary airways express Sox2 alone (Eenjes et al. 2021). Moreover, a narrow Sox2^+^Sox9^+^ transitional area (zone 3) precedes the Sox9^+^ distal tips (zone 4) (Eenjes et al. 2021; Mahoney et al. 2014). In humans, SOX21 similarly marks proximal HFL airways (Eenjes et al. 2021). Our HFL (GW 16–18) analysis revealed that in cartilaginous large airways, TP63^+^ cells were abundant and resided along the basal lamina. In this region, both TP63^+^ and TP63^-^ surface airway cells were SOX21^+^SOX9^-^, whereas evaginating submucosal gland buds were SOX21^+^SOX9^+^ and contained both TP63^+^ and TP63^-^ cells (Figure S1A, top panels). Notably, both surface airway epithelium and submucosal gland buds expressed SOX2 (Figure S1A, bottom panels). In contrast, TP63^+^ airway cells were sparse in non-cartilaginous medium-sized and small airways at this developmental stage. In these regions, we observed SOX2^+^SOX21^+^SOX9^low/+^ and SOX2^+^SOX21^-^SOX9^low/+^ airway cells (Figure S1B), consistent with prior reports of both cell populations in *ex vivo* HFL explants (Eenjes et al. 2*021)*. Together, these findings support categorizing developing human lung epithelium into four molecular zones: (1) SOX2^+^SOX21^+^SOX9^-^ proximal cartilaginous airway cells; (2) SOX2^+^SOX21^+^SOX9^low/+^ intermediate (medium-sized) airway cells; (3) SOX2^+^SOX21^-^SOX9^low/+^ distal (small) airway cells; and (4) SOX2^low/-^SOX9^+^ multipotent distal tip progenitors. Thus, the combinatorial expression of SOX2, SOX21, and SOX9 provided a developmental patterning roadmap to effectively interpret proximal-distal regional identity during lung progenitor differentiation.

Using this roadmap as a guide, we established our strategy with fluorescent reporter hPSCs and subsequently validated it in non**-**reporter lines. Fluorescent reporters enabled purification by cell sorting, eliminating paracrine factors secreted by contaminating cells that could potentially introduce confounding variables – making these lines invaluable for establishing differentiation strategies. At each stage, we examined key lung, airway and basal lineage factors, NKX2-1, SOX2 and TP63, to evaluate differentiation efficiency and isolate basal progenitors or iBCs for molecular and functional analyses. Three dual reporter lines were used. These included two published lines: NKX2-1^GFP^ TP63^tdTomato^ BU3 hiPSC (BU3 NGPT) (Hawkins et al. 2021) and NKX2-1^GFP^ TP63^mCherry^ C17 hiPSC (C17 NGPmC) (Berical et al. 2022), as well as a newly generated NKX2-1^GFP^ SOX2^mRUBY3^ dual reporter in our previously published human embryonic stem cell (hESC) line RUES2 (R2 NGSmR). The R2 NGSmR line was created using our published NKX2-1-GFP construct (Hawkins et al. 2017) and by generating a SOX2-2A-mRUBY3 knock-in reporter at the endogenous SOX2 locus (Figure S1C). Non**-**reporter lines used for validation included BU3 hiPSC and RUES2 hESC.

### Deriving lung progenitors from hPSCs

In Stage 1, lung progenitors were generated using our previously published protocol (Huang et al. 2014; Huang et al. 2015) with modifications optimized for airway differentiation (Figure 1B). Briefly, we previously demonstrated that WNT, BMP and RA signaling are indispensable for lung progenitor specification from hPSCs, whereas FGF10 and FGF7 are dispensable for specification but promote cell proliferation (Huang et al. 2014). Later studies reported that FGF7 potentially supports alveolar differentiation, while FGF10 is involved in airway and basal development (Jacob et al. 2017). Based on these findings, we activated WNT signaling using the pharmacological agonist CHIR99021 (Tocris, hereafter, CHIR), together with BMP, RA and FGF10 signaling to induce lung specification. The resulting day (D) 15 lung progenitors resembled the characteristics of early stage human lung buds (Xu et al. 2016; Nikolic et al. 2017) and expressed NKX2-1, SOX2 and SOX9, as examined by immunofluorescence (IF) (Figure 1C). Notably, D15 cells showed sporadic SOX21 expression (Figure 1D).

### NOGGIN-BMP signaling gradient systematically regulates SOX2, SOX21, and SOX9 expression whereas WNT activation maintains lung fate in hPSC-derived lung progenitors

Stage 2 is critical for proximal-distal regional patterning and NKX2-1 fate maintenance in lung progenitors. To demonstrate the role of NOGGIN-BMP signaling in proximal-distal patterning during Stage 2, sorted D15 NKX2-1^GFP+^ (hereafter, GFP^+^) lung progenitors were embedded in Matrigel droplets for three–dimensional (3D) culture in conditions with serum-free medium containing a NOGGIN-BMP signaling gradient: *a*) proximal, continuous BMP inhibition through its biological inhibitor NOGGIN; *b*) intermediate, endogenous BMPs; and *c*) distal, BMP activation using BMP4 (Figures 1E and S1D). At D30, under proximal conditions, NOGGIN efficiently inhibited SOX9 and the undesired alveolar fate, yielding *<u>proximal airway progenitors</u>* expressing NKX2-1, SOX2 and SOX21 but not SOX9 (Figure 1F-G). Under the intermediate condition, endogenous BMPs promoted *<u>intermediate multipotent progenitors</u>* expressing NKX2-1, SOX2, SOX21 and SOX9. Under the distal condition, BMP4 suppressed SOX21, the proximal airway-associated transcription factor, and promoted *<u>distal multipotent progenitors</u>* that express NKX2-1, SOX2, and SOX9 but lacked SOX21. Under proximal conditions, reducing NOGGIN dose by half (medium NOG, designated as proximal 2) markedly increased SOX21^+^SOX9^+^ cells (17.5 ± 2.8%) compared with high NOG (5.9 ± 1.4%, hereafter proximal 1) (Figure S1E), suggesting that high NOGGIN is required to effectively inhibit endogenous BMPs. Notably, cells under the intermediate condition were SOX21^low^SOX9^+^ or SOX21^+^SOX9^low^ (Figure 1G middle panel, insets). Overall, SOX2, SOX21 and SOX9 expression under our culture conditions was consistent with patterns observed in developing human fetal lungs (Figure S1A-B). Under proximal conditions, we detected increased SOX2^mRUBY3^ reporter protein intensity compared with the intermediate condition, whereas under the distal condition BMP4 suppressed its expression (Figure 1F). The mRNA expression levels of SOX2, SOX21 and SOX9 (Figure 1H) were consistent with their protein levels (Figure 1G). Lower NKX2-1 and higher SOX2 mRNA levels under proximal conditions were consistent with developmental patterning in mouse and human fetal lungs, where proximal airway regions show higher SOX2 and distal regions exhibit a stronger NKX2-1-associated program (Maeda, Dave, and Whitsett 2007; Lim et al. 2023; Cao et al. 2023). Together, these results suggest that NOGGIN-BMP signaling systematically patterns lung progenitors along the proximal-distal axis.

In Stage 2, across all conditions, we continued applying FGF10 signaling for its role in airway basal development and cell growth, as well as CHIR (WNT activation) for its critical role in lung fate maintenance. We previously reported that embryonic day (E) 11 mouse lung and hPSC-derived D12–15 lung progenitors require tonic WNT signaling for NKX2-1 maintenance (Ostrin et al. 2018). Here we show that NKX2-1 maintenance remained WNT-dependent during D15–30 in three hPSC lines (Figure S1F). We compared conditions with and without WNT activation using CHIR. High CHIR maintained NKX2-1^GFP^ expression in most cells, while adding NOGGIN or BMP4 reduced NKX2-1 maintenance; without CHIR, most cells lost GFP expression (Figure S1F-G). At this stage, TP63 reporter-positive cells were sparse under high CHIR conditions (Figure S1G), consistent with prior reports that airway basal-cell specification requires WNT downregulation (Hou et al. 2019; Hashimoto et al. 2012; Mucenski et al. 2003; Shu et al. 2005; McCauley et al. 2017; Hawkins et al. 2021).

Chromatin immunoprecipitation sequencing (ChIP-seq) confirmed that transcription factor 4 (TCF4), a WNT/CTNNB1 effector, binds *NKX2-1* in D17 lung progenitors (Figure S1H), suggesting that WNT signaling directly regulates *NKX2-1* and that WNT withdrawal impacts NKX2-1 expression. Together, these data support a differentiation strategy in which condition-specific NOGGIN–BMP modulation establishes proximal–distal regional identity, whereas WNT signaling is temporally tapered across Stages 2–4 to promote basal specification while preserving NKX2-1+ lung fate (Figure 1A).

Specifically, Stage 2 high WNT maintained NKX2-1^+^ lung progenitor fate, Stage 3 low WNT promoted TP63^+^ specification while maintaining NKX2-1 expression, and Stage 4 WNT withdrawal/endogenous WNT supported basal maturation. This differs from prior published schemes, in which WNT is withdrawn immediately after NKX2-1^+^ lung progenitor specification at D15 (Hawkins et al. 2021; Wang et al. 2023). Notably, TCF4 also bound *SOX2* (Figure S1H).

Under the distal condition, Stage 2 was extended to D45, providing an additional 15 days of high-level WNT activation required to stabilize lung fate before WNT reduction and basal specification. Thus, compared with proximal or intermediate progenitors, distal progenitors required more prolonged Stage 2 WNT activation to maintain lung fate and preserve subsequent basal cell competence through Stages 3–4.

### Generation of region-specific iBCs using a NOGGIN-BMP gradient and tapered WNT signaling

In Stage 3, we applied low CHIR to reduce WNT activity (Figure 2A), thereby promoting airway fate commitment and basal specification. At the end of Stage 3 (D45 for proximal and intermediate conditions; D60 for the distal condition), the combination of low CHIR and high NOGGIN (proximal 1) rapidly specified most cells into putative *<u>proximal basal progenitors</u>*, as defined in Figure 1A (Figure 2B). Notably, the proximal 2 condition (low-CHIR, medium-NOGGIN) also efficiently generated NKX2-1^+^/TP63^+^ basal progenitors (Figure S2B), which contained a higher proportion of SOX21^+^SOX9^+^ cells at D30 than proximal 1 progenitors (Figures 1G and S1E). Under the low-CHIR-only condition (endogenous BMPs), a mixture of putative *<u>intermediate basal progenitors</u>* (TP63^+^) and *<u>airway progenitors</u>* (TP63^-^) was generated (Figure S2B). The low-CHIR plus BMP4 condition generated a mixture of putative *<u>distal basal progenitors</u>* (TP63^+^) and *<u>airway cells</u>* (TP63^-^) (Figure 2B).

**Figure 2.**
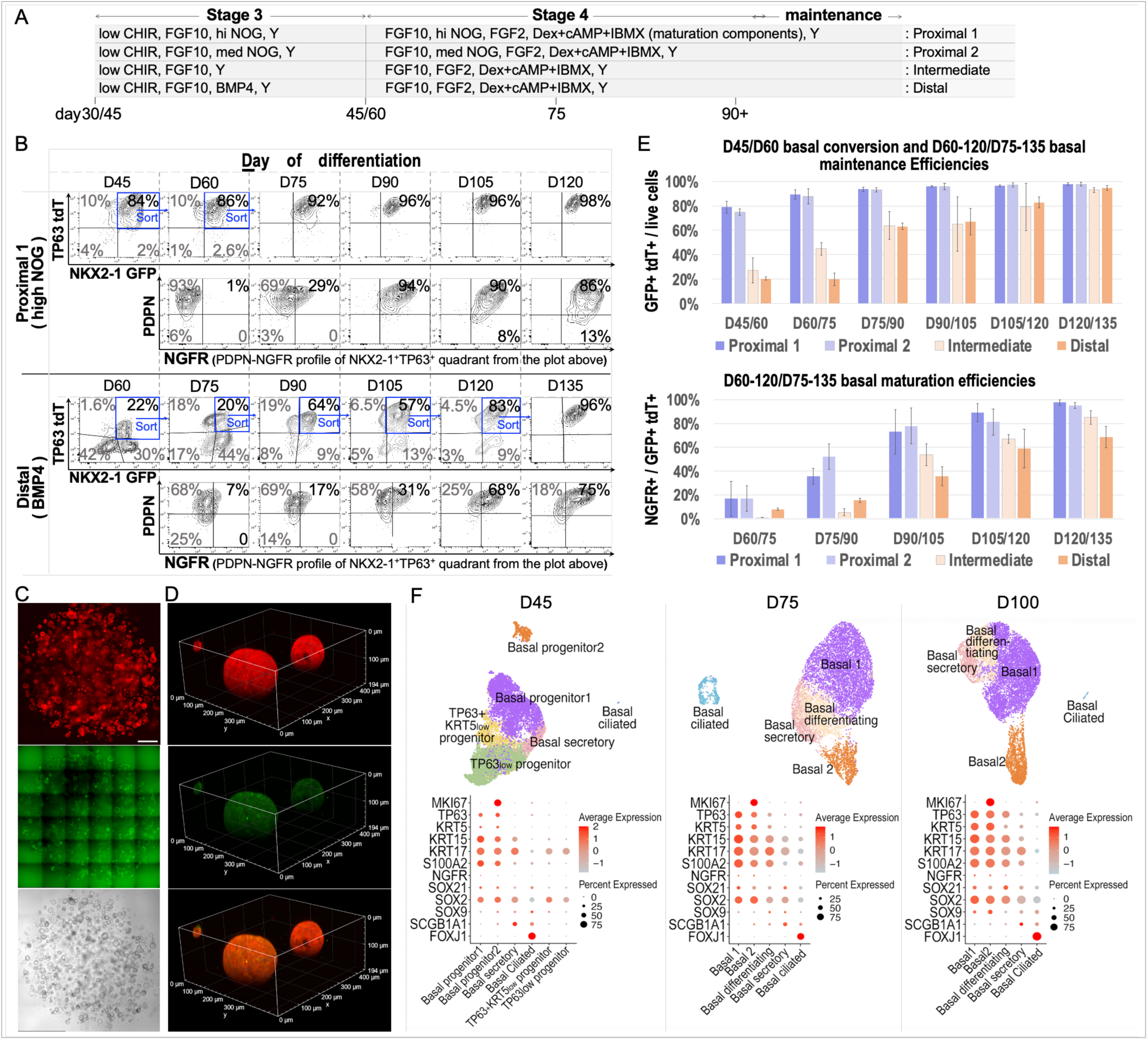
Generation of region-specific iBCs using a NOGGIN-BMP gradient and tapered WNT signaling. (**A**) Schematic of the stages 3–4 culture protocol from days 30/45 to 90+ (D30–90+ for proximal and intermediate conditions; D45–105+ for the distal condition). Dex, dexamethasone; cAMP, cyclic AMP; IBMX, isobutylmethylxanthine. Notably, under the distal condition, basal progenitor specification is delayed and begins at D45. (**B**) Basal progenitor specification, maintenance, and maturation under the indicated experimental conditions. Basal progenitor specification efficiency was measured as the percentage of NKX2-1^GFP+^ TP63^tdT+^ cells at the end of stage 3 (D45 for proximal and intermediate conditions; D60 for the distal condition); basal maintenance during stage 4 was assessed by the percentage of cells remaining GFP^+^tdT^+^ at each time point; basal maturation was measured by NGFR expression in GFP^+^tdT^+^ cells. *n* = 3 for each hPSC line, a representative experiment from BU3 NGPT is shown for proximal 1 and distal conditions. Notably, some proximal NGFR^+^ iBCs lost PDPN expression at D105-120. (**C**) 6 x 6 tiled 10x images of D60 proximal 1 basal progenitors embedded in a Matrigel droplet. Scale bar: 1mm. (**D**) Confocal image of two D60 basal progenitor organoids. (**E**) Average efficiencies of basal progenitor specification, maintenance, and maturation for each of the proximal-distal condition across three experiments from BU3 NGPT line. (**F**) Molecular signatures of basal (*TP63, KRT5, KRT15, KRT17, S100A2, NGFR*), proliferative (*MKI67*), proximal-distal patterning (*SOX2, SOX21, SOX9*), secretory (*SCGB1A1*), and ciliated (*FOXJ1*) markers in D45, D75 and D100 proximal 1 cells from BU3 NGPT assessed by scRNA-seq. UMAPs (*top*) display the identified clusters, and corresponding dot plots (*bottom*) show expression of the indicated markers across clusters.

In some experiments, TP63⁻ cells were sorted and cultured for an additional 15 days in Stage 3 basal specification medium. After this extended Stage 3 period (D45–60 for proximal and intermediate conditions; D60–75 for the distal condition), only a small fraction of cells expressed TP63, while most retained an airway and/or secretory progenitor phenotype. These findings suggest that ∼15 days of Stage 3 culture were generally sufficient for most cells to initiate TP63 expression. However, because low CHIR in Stage 3 also helped stabilize lung fate, extending basal specification for an additional 15 days may be beneficial or necessary in some hPSC lines before transition to basal maturation.

In Stage 4, the putative basal progenitors (NKX2-1^GFP+^ TP63^tdTomato+^ or NKX2-1^GFP+^ TP63^mCherry+^; hereafter, GFP^+^tdT^+^ or GFP^+^mCh^+^) from each condition were purified for maturation without CHIR, mirroring findings in mouse studies (Hou et al. 2019; Hashimoto et al. 2012; Mucenski et al. 2003; Shu et al. 2005). In some hPSC lines, extending Stage 3 improved lung fate stability after transition to Stage 4. For example, in the BU3 NGPT line, switching to Stage 4 at D60 rather than D45 better preserved NKX2-1 GFP expression in proximal cells at D75 and later time points in some experiments. Under proximal conditions, NOGGIN was required because basal progenitors continuously produced endogenous BMPs (Figure S2C). Under the distal condition, BMP4 was withdrawn to avoid excessive suppression of the airway-associated transcription factor SOX2 and to permit basal maturation. Under proximal conditions (1 and 2), most cells maintained NKX2-1 and TP63 reporter protein expression from D60 onward, as assessed by FACS and imaging of organoids (Figure 2B-D). The expression of the mature basal surface marker NGFR (Rock et al. 2009) increased progressively, and by approximately D90, most cells expressed NGFR (Figures 2B, 2E and S2B), which we define as putative *<u>proximal basal cells</u>* (Figure 1A), or <u>proximal iBCs</u>. Under intermediate and distal conditions, a proportion of the initially sorted TP63^+^ cells turned off TP63 and assumed a differentiating airway progenitor phenotype, with TP63 fate stabilizing around D75–90 (Figures 2B, 2E and S2B). We define the stabilized TP63^+^ populations as putative *<u>intermediate and distal airway basal cells</u>*, or <u>intermediate and distal iBCs</u>. This phenomenon is consistent with findings in the developing mouse lung, where only a subset of TP63^+^ cells commits to basal fate postnatally (Yang et al. 2018). The lower basal specification and maintenance observed under intermediate and distal conditions also align with the lower abundance of airway basal cells along the proximal-distal axis of the airway tree.

To address the important question of whether the gradual increase and maintenance of NGFR^+^ cells from D60 to D120 was due to a relative proliferation advantage of NGFR^+^ over NGFR^-^ cells, we sorted NGFR^+^ and NGFR^-^ cells from proximal 1, proximal 2, intermediate, and distal conditions during D75–120 and cultured them separately in 3D droplets. We checked the NGFR profiles every 15 days. The outcomes were consistent across all four conditions; here, we report representative results from the proximal 1 condition. The data consistently showed that both NGFR⁺ and NGFR⁻ populations reached similar profiles 10–14 days after sorting. At D75, approximately 40% of the cells in both the D60 sorted NGFR⁺ and NGFR⁻ populations expressed NGFR (Figure S2E). By D90, 95% of the D75 sorted NGFR⁻ cells had begun expressing NGFR within two weeks (Figure S2E). Conversely, about 5% of the D75 sorted NGFR⁺ cells lost NGFR expression (Figure S2E). We then performed a second round of NGFR^+^ vs. NGFR^-^ sorting and embedded each of the four populations separately in 3D droplets. Fifteen days later, each of the four populations showed similar NGFR profiles, with the majority of cells expressing NGFR (Figure S2E). These results indicate that region-specific iBCs retained plasticity, with NGFR^+^ cells able to revert to an NGFR^-^ progenitor state and NGFR^-^ cells able to reacquire NGFR expression. Importantly, region-specific iBCs could be expanded in our defined basal medium for at least six months, maintained the NGFR profile, and exhibited robust differentiation potential.

We next translated the reporter-informed framework into a surface marker-based sorting strategy for generating proximal, intermediate, and distal iBCs from non-reporter hPSCs. Using reporter lines, we identified surface markers for isolating NKX2-1–expressing lung progenitors (EpCAM^+^carboxypeptidase M [CPM]^hi/+^) during Stages 1–2, enriching putative TP63⁺ basal progenitors (EpCAM^+^F3^+^PDPN^+^) during Stages 3–4, and purifying EpCAM⁺NGFR⁺ iBCs during Stage 4. Consistent with prior reports (Gotoh et al. 2014), CPM^hi^ (D15) and CPM⁺ (D30) populations closely correlated with NKX2-1^GFP^⁺ cells under high CHIR during Stages 1–2 (Figure S3A–B). Lung fate, as indicated by CPM expression, was largely stabilized by the end of Stage 2 (D30 under proximal/intermediate conditions; D45 under the distal condition) (Figure S3C). By the end of Stage 2, the CPM⁻ fraction showed reduced forward scatter (FSC), consistent with dissociation-induced stress, and failed to survive upon further culture. These cells could be largely excluded by propidium iodide (PI) staining during sorting, although PI was omitted for cells intended for downstream culture to avoid toxicity. Following CHIR reduction from Stage 3 onward, CPM expression was no longer specific to GFP⁺ cells (Figure S3B, *top right*). However, NGFR⁺ cells arose exclusively from the GFP⁺ fraction (Figure S3B, *top right*), indicating that loss of CPM specificity does not compromise downstream purification. Under proximal conditions, basal specification efficiency was high, and GFP loss was primarily associated with dissociation-induced stress. Under intermediate and distal conditions, TP63⁺ cells similarly arose predominantly from the GFP⁺ fraction (Figures 2B and S2B). Thus, enrichment of basal progenitors during Stages 3–4 followed by NGFR-based sorting was sufficient to isolate iBCs.

F3 and EGFR, previously reported to be enriched in fetal lung basal populations (Miller et al. 2020), were relatively low and uniformly expressed at D30 (Figure S3C) but increased and became closely correlated with each other following WNT downregulation during Stages 3–4, with F3 exhibiting broader distribution (Figure S3D, Stage 3 panels). PDPN has also been reported in basal cells together with NGFR (Rock et al. 2009). Although not specific to basal progenitors, PDPN enriched TP63^tdT^⁺ populations across all conditions at the end of Stage 3 basal specification (D45 for proximal/intermediate conditions; D60 for the distal condition; Figure S3B, bottom), and showed the broadest distribution overall (Figure S3D, Stage 3 panels). Notably, NGFR⁺ cells were detected predominantly within the F3⁺ and PDPN^hi^ fractions (Figure S3D; bottom panels).

Accordingly, we sorted F3⁺PDPN⁺ populations across the Stage 3-to-Stage 4 transition (D45–60 for proximal/intermediate conditions; D60–75 for the distal condition) to enrich TP63⁺ progenitors. Under proximal conditions, these populations were stably maintained beyond D60, and additional sorting was not required. In contrast, under intermediate and distal conditions, F3⁺PDPN⁺ cells partially lost marker expression after sorting (Figure S3D), consistent with the loss of TP63 reporter expression observed in a subset of tdT^+^ cells in reporter lines (Figures 2B, S2B). Under these conditions, we applied a more stringent sorting strategy, involving isolation of F3^hi^PDPN^hi^ populations during early Stage 4 basal maturation (D60 for the intermediate condition; D75 for the distal condition), followed by separation of PDPN^hi^ and PDPN^+^ populations 15 days later (D75 and D90, respectively) for further culture (Figure S3D, red oval gates). This improved maintenance of basal identity, limited overproliferation of airway progenitors that would otherwise outcompete basal progenitors, and promoted NGFR expression, consistent with enrichment of NGFR potential in the PDPN^hi^ fraction (Figures 2B, S2B, and S3D; bottom panels of each condition). Beyond these time points, F3⁺ sorting was used as needed to enrich basal populations because reporter data showed persistence of a small fraction (∼5%) of PDPN⁻NGFR⁻ basal progenitors at D90+ and loss of PDPN expression in some NGFR⁺ iBCs beyond D100–120 (Figures 2B, S2B, and S2E). In addition, as supported by reporter line experiments (Figure S2E), separating NGFR⁺ and NGFR⁻ populations for further culture (Figure S3D, red rectangular gates) promoted NGFR expression and enhanced iBC maturation, particularly under the distal condition. Consistent with reporter line results, intermediate and distal conditions showed lower basal specification and maintenance efficiency than proximal conditions. Although this pattern aligns with *in vivo* airway basal cell distribution, these results also indicate that additional signaling regulation may be needed to improve basal specification and maintenance under the distal condition. Given the robust generation of proximal iBCs and the lack of prior methods for directed proximal iBC differentiation, we focused our in-depth characterization on this population.

We next defined the molecular phenotypes of putative proximal basal progenitors/iBCs by single-cell analysis at D45, D75, and D100. For D45 sample preparation, NKX2-1^GFP+^TP63^tdT+^ single live cells maintained in basal specification medium in 3D Matrigel droplets under the proximal 1 condition were sorted for analysis. These cells lacked NGFR expression by antibody staining and FACS. D75 cells were prepared by sorting D60 GFP^+^tdT^+^ proximal 1 cells and culturing them for an additional 15 days in defined basal maturation/maintenance medium. At D75, ∼92% of cells remained GFP^+^tdT^+^, and ∼30% expressed NGFR protein. For D100, GFP^+^tdT^+^NGFR^+^ proximal 1 and proximal 2 cells were sorted at D90 and cultured for an additional 10 days in defined basal maintenance medium. At D100, ∼96% of cells remained GFP^+^tdT^+^, and 85%–95% expressed NGFR. Quantitative kinetics are shown in Figures 2B, 2E and S2B. For each time point, 10,000 sorted single live cells from proximal 1 were processed for 10x Genomics scRNA-seq, and 10,000 sorted single live cells from proximal 2 were processed for 10x Genomics sc-multiome analysis; only the gene expression component of the sc-multiome dataset was analyzed in this study. Dot plots (Figure 2F) showed a progressive increase in the fraction of proximal 1 cells expressing canonical basal and airway marker transcripts (*TP63, KRT5, KRT15, KRT17, S100A2, SOX2*) from D45 to D100 at ∼40% sequencing depth, with most D100 cells expressing basal markers. D100 proximal 1 and proximal 2 cells exhibited highly similar expression patterns (Figures 2F, S2D). These cells segregated into two major clusters, basal progenitor 1/basal 1 and basal progenitor 2/basal 2, the latter comprising *MKI67^+^* cycling cells. *NGFR* (mature BC marker) and *SOX21* (proximal patterning factor) were underrepresented at the transcript level relative to their protein abundance detected by antibody staining (Figures 2F, S2D vs. Figures 1G, 2B, and S2B). Sporadic cells expressed *SOX9* transcripts, consistent with protein expression. The D45 sample contained *TP63^low^ KRT5^low^* progenitor clusters, suggestive of developing basal progenitors with low transcript abundance that were likely underrepresented because of sequencing depth, despite clear TP63 reporter detection by FACS. scRNA-seq also identified a small “basal differentiating” population with reduced *TP63* and *KRT5* at D75 and D100, as well as minor TP63^low^ subsets co-expressing secretoryglobulin family 1A member 1 (*SCGB1A1*; “basal-secretory”) or forkhead box *(FOX) J1* (“basal-ciliated”) across all time points (Figures 2F and S2D). Together, these data support efficient generation and maintenance of proximal iBCs using this protocol.

### Proximal iBCs are fully functional *in vitro* and *in vivo*

After defining the molecular phenotypes of proximal iBCs, we functionally characterized them *in vitro* and *in vivo*. To assess their differentiation potential *in vitro*, D90–130 proximal iBCs were seeded onto Transwell inserts and cultured in Pneuma-Cult air-liquid interface (ALI) medium (STEMCELL Technologies) for 4 weeks (Figure 3A). Resulting ALI cultures contained cells expressing FOXI1 and ASCL1, the transcriptional drivers of ionocyte and PNEC lineages, respectively (Figure 3B–F). By whole-mount IF, a subset of FOXI1⁺ cells co-expressed Barttin CLCNK Type Accessory Subunit Beta (BSND) (Figure 3B), recently reported as one of the three ionocyte subtypes in humans and ferrets (Yuan et al. 2023). In transverse paraffin sections, most FOXI1⁺ cells localized near the apical surface of the differentiation culture, while a few FOXI1⁺BSND⁻ cells were detected near the basal membrane (Figure 3C), suggestive of a progenitor state as recently reported (Shah et al. 2025). Notably, most ionocytes expressed CFTR on the apical membrane (Figure 3D). Overall, proximal 1 iBCs showed robust ionocyte differentiation (Figure 3B–C). Whole-transwell tiles across lines are shown in Figures S4–S5. FOXI1⁺ cells constituted 1.2–3.1% of the cultures based on counts of ∼1,000 cells on transverse sections, and FOXI1⁺BSND⁺ cells comprised 33–65% of FOXI1⁺ cells (Figure 3E). Ionocyte subtype composition in proximal iBC-ALI cultures will be examined in future work. Because recent studies suggest airway rare cell types may arise from shared progenitors (Yuan et al. 2023; Shah et al. 2025), we assessed FOXI1/ASCL1 co-expression: whole-mount IF revealed largely mutually exclusive populations with only sporadic double-positive cells (two FOXI1⁺/^low^ASCL1^low^ cells in the field shown; Figure 3F), indicating lineage commitment and segregation of ionocyte and PNEC cell fate at this stage. Tuft cells, although enriched proximally *in vivo* (Montoro et al. 2018; Plasschaert et al. 2018), were not detected in our ALI cultures, consistent with a recent report (Shah et al. 2025) showing tuft cells are rare *in vivo* at steady state and in ALI cultures, but increase upon specific cytokine stimulation. Lastly, major airway lineages were present in our ALI cultures, including secretory (SCGB1A1, mucin 5AC [MUC5AC]) and multiciliated (acetylated α-tubulin [ACT]) cells (Figure S6). The basal–apical distribution of airway lineage markers is shown by IF on transverse paraffin sections (Figure 3G).

**Figure 3.**
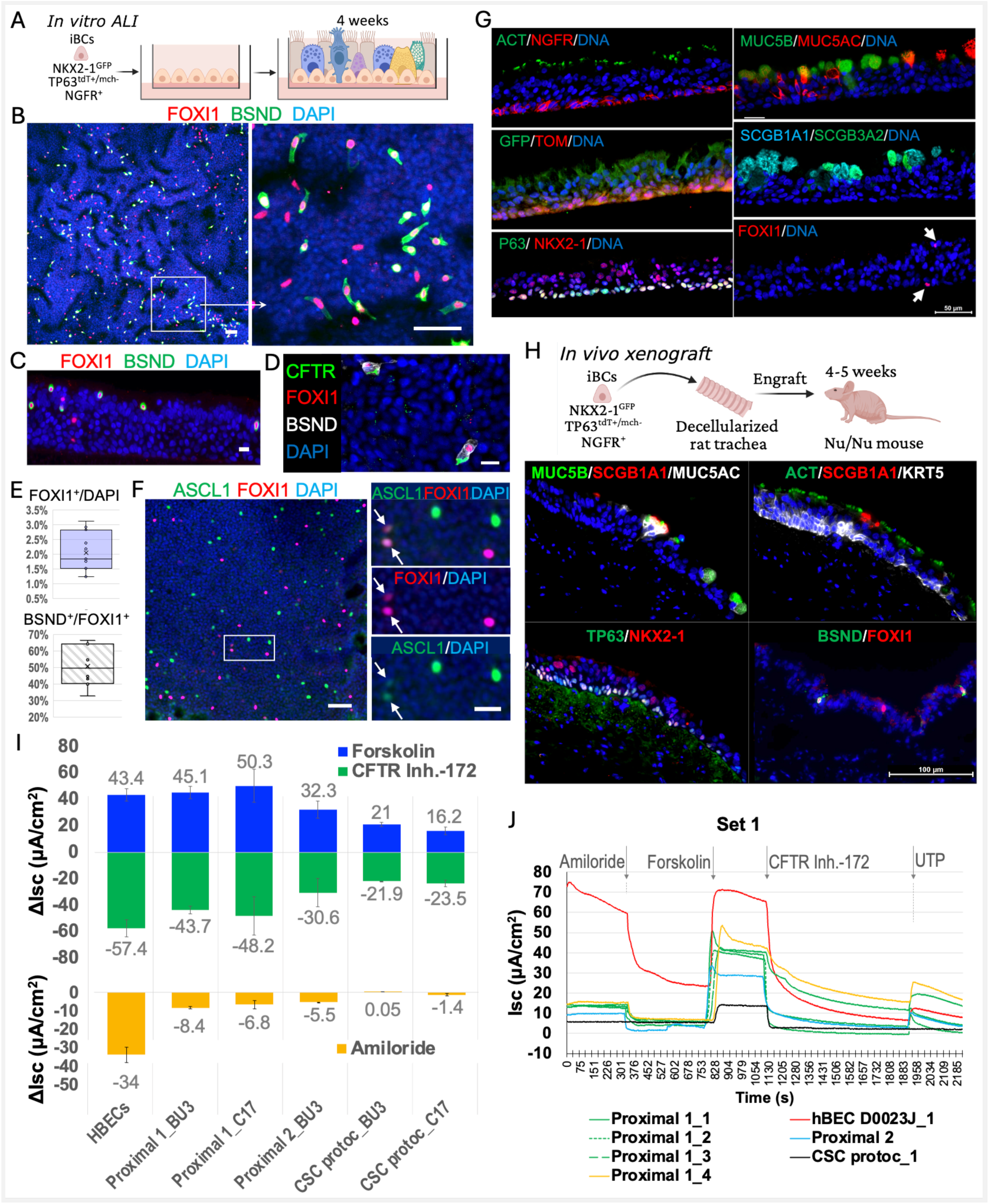
Functional characterization of proximal iBCs *in vitro* and *in vivo*. D90–130 proximal iBCs were differentiated in ALI medium for 4 weeks *in vitro* (A) or in a tracheal xenograft model for 4 weeks *in vivo* (H). (**A**) Schematic of the ALI differentiation protocol. (**B–F)** Characterization of ionocytes and PNECs in proximal 1 iBC-ALI cultures by FOXI1/BSND whole mount (B) and paraffin section (C) staining, CFTR/FOXI1/BSND (D), and FOXI1/ASCL1 (F) whole mount staining. Scale bar: 50μm (B and F, *left*) and 20μm (C, D, and F, *right*). E: quantification of FOXI1^+^ and BSND^+^/FOXI1^+^ cell populations. *n* = 3 for each hPSC line; representative images from hiPSC line BU3NGPT. (**G–H**) Proximal iBCs exhibited multipotent differentiation *in vitro* in ALI cultures and *in vivo* in tracheal xenografts. Cell type-specific markers were detected with indicated antibodies on paraffin sections of ALI cultures (G) or tracheal xenografts (H). Scale bars: 50μm. n=3 for each hPSC line; representative images from hiPSC line BU3NGPT are shown. (**I–J**) Electrophysiological measurements of proximal iBC- and HBEC-derived ALI cultures by Ussing chamber assay. (I) Average ΔIsc values for each condition from HBECs and the BU3 NGPT & C17 NGPmC lines. Two experiments per line and 2-4 inserts tested per experiment. (J) Representative tracings of individual Transwell inserts showing transepithelial chloride transport under the indicated conditions in BU3 NGPT cultures. CSC protoc. refers to the CSC protocol described in our previous publication (Hawkins and Suzuki *et al*, *Cell Stem Cell*, 2021).

The *in vivo* differentiation capacity of proximal iBCs (D90–130) was assessed using the tracheal xenograft model originally developed for adult human bronchial epithelial cells (HBECs) and subsequently applied in multiple studies (Everitt et al. 1989; Engelhardt et al. 1993; Goldman, Yang, and Wilson 1995; Dupuit et al. 2000; Filali et al. 2002; Hawkins et al. 2021), including the initial evaluation of iBCs generated using our previously published method (Hawkins and Suzuki *et al*, <u>C</u>ell <u>S</u>tem <u>C</u>ell 2021, hereafter termed the CSC protocol). In this model, fully decellularized rat tracheas are fitted with tubing to generate open-ended xenografts that allow luminal air exposure, while cells seeded on the denuded trachea scaffold are nourished by the host systemic circulation after transplantation into mice, thereby recapitulating the *in vivo* environment. Xenografts were prepared, seeded with proximal iBCs and implanted subcutaneously into the flanks of *Foxn1^nu^* immunodeficient mice as previously described (Filali et al. 2002) with minor modifications. At four to five weeks post-engraftment, immunofluorescence of xenograft sections showed differentiation of seeded proximal iBCs into multiple airway lineages, including ionocytes (FOXI1/BSND) (Figure 3H). Basal cells (TP63/KRT5) localized along the basement membrane; multiciliated cells (ACT) oriented toward the tracheal lumen; and secretory-cell markers (SCGB1A1/MUC5AC/MUC5B) exhibited apical polarization. Overall, this model demonstrated that transplanted proximal iBCs can generate a stratified airway epithelium resembling *in vivo* airways. The engraftment and reconstitution capacity of our region-specific iBCs in an airway injury transplantation model (Ma et al. 2023) will be evaluated in future studies.

Next, we evaluated CFTR anion channel activity in proximal iBC-derived ALI cultures using an Ussing chamber assay. As a control, we used ALI cultures derived from HBECs, which are culture-expanded adult human basal cells isolated from the proximal airway region. Additionally, we analyzed ALI cultures derived from iBCs differentiated using our previously published CSC protocol (Hawkins et al. 2021) side by side. The results showed that CFTR activities (Forskolin induced and CFTRinh-172 inhibited) of ALI cultures derived from proximal 1 iBCs were comparable to those of HBEC-derived ALI cultures (Figure 3I, top). Importantly, these ALI cultures also exhibited sodium channel activity (Figure 3I, bottom). In comparison, ALI cultures derived using the CSC protocol exhibited lower CFTR activity, and sodium channel activity was largely absent (Figure 3I). The tracings of individual Transwell inserts for transepithelial chloride transport in response to Forskolin stimulation followed by the inhibitor CFTRinh-172 treatment are shown in Figure 3J. ΔIsc was calculated by subtracting the pre-stimulation/treatment Isc values from the post-Forskolin/CFTRinh-172 values.

### Region-specific iBC-ALI cultures recapitulate the molecular signatures and phenotypes of large, medium, and small human airways

We also systematically examined the differentiation potential of region-specific iBCs. Specifically, D90–130 region-specific iBCs and HBECs were plated on Transwell inserts for differentiation in ALI medium (Pneuma-Cult ALI) for 4 weeks. In some experiments, ALI cultures derived from iBCs differentiated using the CSC protocol were included as controls. We performed scRNA-seq on ALI cultures derived from proximal 1, proximal 2, and intermediate iBCs, as well as HBECs, to compare the molecular signatures of the differentiated cell types. Using established molecular signatures for each airway cell type, we identified clusters of major cell types, including basal, cycling basal, basal transitional, secretory, secretory transitional, ciliated, *FOXN4^+^*(deutosomal), and ciliated transitional cells in all conditions (Figure 4A-D). Clusters of rare cell types, such as ionocytes and PNECs, were present in HBEC- and proximal 1 iBC-derived ALI cultures (Figures 4A-B and S7A), while in proximal 2 iBC-ALI, ionocytes and PNECs clustered together (Figures 4C and S7B). In intermediate iBC-derived ALI cultures, few ionocytes and PNECs were captured by scRNA-seq, and they clustered with basal transitional cells (Figures 4D and S7C). Notably, in HBEC- and proximal iBC-ALI cultures, the relative expression of *CFTR* was higher in individual ionocytes than in secretory cells, consistent with published findings in humans, mice, and ferrets (Montoro et al. 2018; Okuda et al. 2020; Plasschaert et al. 2018; Travaglini et al. 2020; Yuan et al. 2023). We then integrated the HBEC-ALI dataset with three independent proximal 1 iBC-ALI datasets, including one derived from C17 NGPmC and two from BU3 NGPT, using Seurat’s unsupervised integration. The results showed that clusters were driven by cell type rather than dataset; each cluster was well represented in all four individual datasets (Figure 4E), indicating comparable lineage composition and differentiation potential between HBECs and proximal 1 iBCs. The top 5–25 differentially expressed genes (DEGs) per cluster for the proximal 1 condition are shown in Figure 4F. Each cluster exhibited characteristic transcriptional drivers and phenotypic markers defining cell-type-specific populations (Figure 4F). Feature plots for exemplar DEGs are provided in Figure S7A, including *TP63, KRT5*, and *KRT15* (basal); *KRT13* (basal transitional; top DEG), *SCGB3A1* (secretory, third-highest DEG); *FOXJ1* (multiciliated); *FOXI1* and *ASCL3* (ionocyte; sixth- and fourth-highest DEGs); *ASCL1* and *CALCA* (PNEC, seventh- and second-highest DEGs).

**Figure 4.**
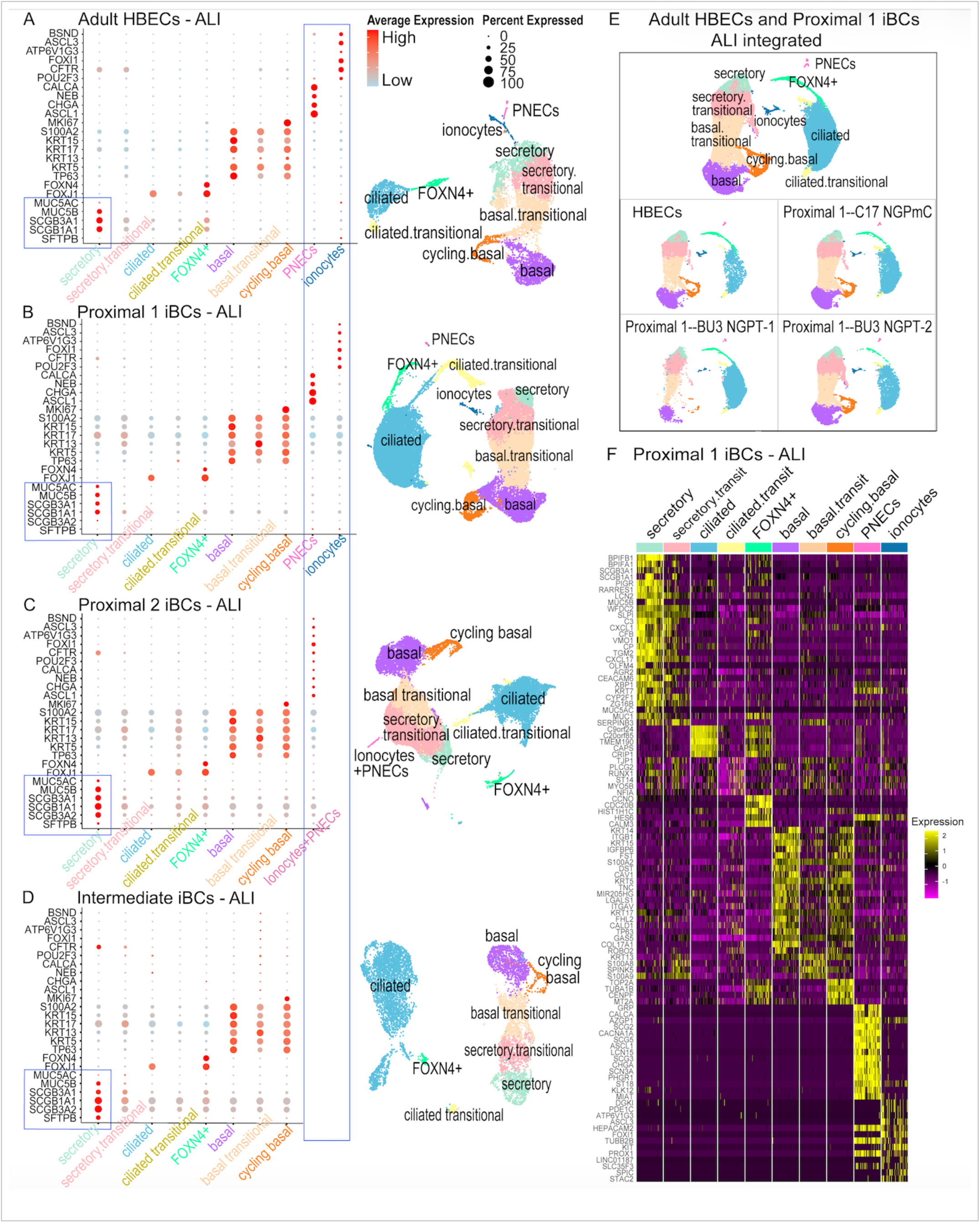
Region-specific iBC-ALI cultures exhibit molecular signatures of differentiated cell types characteristic of human large and medium airways. (**A–D**) HBECs and iBCs were differentiated according to Figure 3A, and live single cells from each culture condition were sorted by flow cytometry for 10x Genomics scRNA-seq. Dot plots (*left*) show expression of selected markers in each condition for the cell types indicated in the UMAPs (*right*). Signatures of secretory cells, ionocytes and PNECs in each condition are highlighted by blue rectangles. A, ∼10,000 cells from one dataset; B, ∼30,000 cells from three integrated independent datasets; C, ∼20,000 cells from two integrated independent datasets; D, ∼10,000 cells from one dataset. (**E**) UMAPs of integrated HBEC-ALI and proximal 1 iBC-ALI cultures, as well as of each individual sample. (**F**) Heatmap showing the top 5**–**25 DEGs for each cluster in integrated proximal 1 iBC-ALI cultures. For secretory cells, the top 25 DEGs are shown; for basal cells, the top 20 DEGs are shown; for PNECs and ionocytes, the top 15 DEGs are shown; and for all the other clusters, the top 5 DEGs are shown.

As detailed earlier, *ex vivo*–cultured human proximal and distal airway BCs differentiate into region-appropriate progeny, and secretory cells from proximal, intermediate, and distal human airways exhibit region-specific phenotypes (Kadur Lakshminarasimha Murthy et al. 2022; Zhou et al. 2022). Guided by these findings, we examined secretory profiles under each condition (Figure 4A–D) and compared them with published adult human airway regional secretory signatures and spatial patterning (Kadur Lakshminarasimha Murthy et al. 2022). The secretory cells from both HBEC- and proximal 1 iBC-derived ALI cultures exhibited the combinatorial signatures of Zone-1 large airway secretory cells, including *SCGB1A1, SCGB3A1, MUC5AC, MUC5B*, and the absence of *SCGB3A2* and *surfactant protein B (SFTPB)* (Figures 4A-B and 5A). Proximal 2 iBC-differentiated secretory cells mapped to Zone 2, expressing *SCGB1A1, SCGB3A1*, and *MUC5B*, with reduced *MUC5AC* and increased *SCGB3A2* and *SFTPB* expression (Figures 4C and 5A). This pattern was consistent with detection of some SCGB3A2 protein by IF in proximal 2 iBC-derived ALI cultures (Figure 3G). Intermediate iBC-derived secretory cells mapped to Zone 2 and 3. They expressed *SCGB1A1*, *SCGB3A2* and *SFTPB*, lacked *MUC5AC*, and showed reduced *SCGB3A1* and *MUC5B* expression (Figures 4D and 5A). The transcriptional signatures from proximal 1 and intermediate conditions were largely consistent with the protein expression profiles by IF (Figure 5B). In keeping with this, SCGB3A2 protein was nearly absent in proximal 1 iBC-derived ALI cultures, detected at low levels under the intermediate condition, and robustly expressed under the distal condition. Under the distal condition, secretory cells showed robust SCGB3A2 and SFTPB expression, reduced SCGB1A1 expression relative to proximal and intermediate conditions, and sporadic SCGB3A1, MUC5AC, and MUC5B expression (Figure 5B), mapping to Zone 3 adult human secretory cells (Kadur Lakshminarasimha Murthy et al. 2022). Furthermore, we confirmed the mRNA expression of these secretory signatures by qPCR (Figure 5C). Notably, low levels of *MUC5B* and *SCGB3A1* mRNA were detected in the distal iBC-derived secretory cells; however, the overall expression pattern aligns with protein expression observed by IF and further validates the scRNA-seq data.

**Figure 5.**
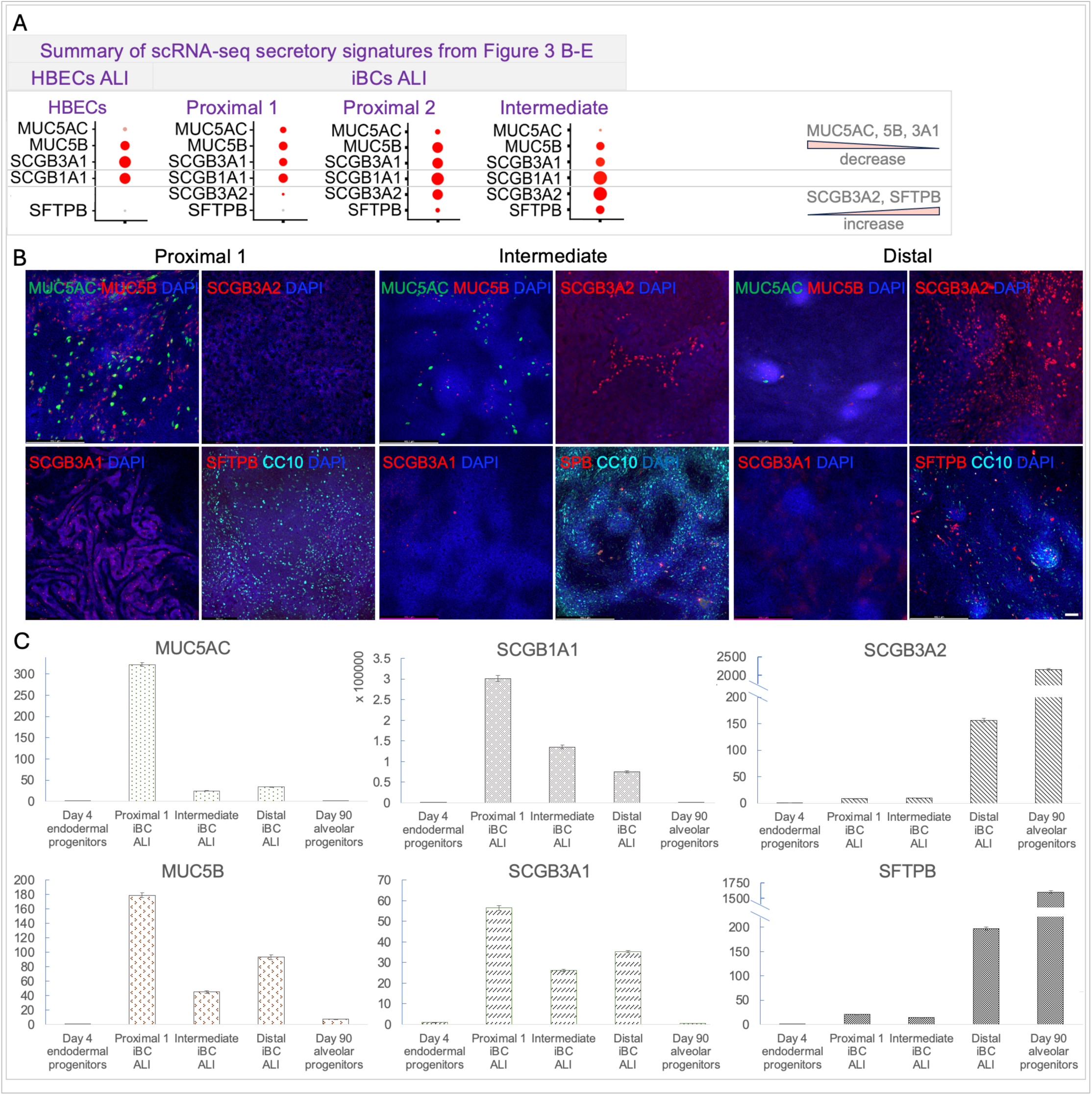
RNA and protein signatures in region-specific iBC-ALI cultures indicate secretory phenotypes mapping to Zones 1–3 of adult human airways. (**A**) summary of the secretory signatures from Figure 4A–D. (**B**) Characterization of MUC5AC, MUC5B, SCGB31A1, SCGB3A1, SCGB3A2 and SFTPB by immunofluorescence in region-specific iBC-ALI cultures differentiated as shown in Figure 3A. *n* = 3 for each hPSC line; a representative experiment from C17 NGPmC is shown. Scale bar: 100μm. (**C**) Verification of *MUC5AC, MUC5B, SCGB31A1, SCGB3A1, SCGB3A2* and *SFTPB* mRNA expression by qPCR under the indicated conditions. Expression levels are shown normalized relative to D15 lung progenitor condition cultured according to Figure 1B, *n* = 3 for each cell line; a representative experiment from C17 NGPmC is shown.

To assess the regional identity of major differentiated airway cell types while accounting for culture-associated effects, we next directly compared basal, secretory, and ciliated-cell signatures from both the proximal 1 iBC and HBEC datasets with those from freshly isolated, tissue-resident cells of human proximal and distal airways. This comparison was important because *in vitro* culture can alter primary-cell phenotypic characteristics, and our adult HBEC positive controls were also differentiated in ALI cultures. Included for analysis were three published adult human proximal-airway scRNA-seq datasets (Okuda et al. 2020; Carraro et al. 2021; Basil et al. 2022), two distal-airway datasets (Basil et al. 2022; Kadur Lakshminarasimha Murthy et al. 2022), and the LUNGMAP meta-dataset (proximal and distal combined) (Guo et al. 2023). Analyses were performed with 4,500 highly variable genes (HVGs) per cell type across datasets – more stringent than the default 2000. Overall, despite culture-associated effects, proximal iBC-ALI and HBEC-ALI differentiation cultures contained basal and secretory cells that best aligned with tissue-resident counterparts in human proximal, but not distal, airways (Table 1). Notably, micro-dissected distal airway samples contained basal and secretory cells exhibiting proximal signatures (Kadur Lakshminarasimha Murthy et al. 2022); their potential origins are described in Table 1 footnote. Accordingly, these cells showed stronger correlation with basal and secretory cells from the proximal 1 iBC and HBEC datasets. By contrast, ciliated-cell correlation coefficients were similar across regions, indicating comparable overall HVG transcriptional profiles for ciliated cells in proximal and distal airways by scRNA-seq (Table 1). Regional identity of ciliated cells may therefore be better assessed by morphological and functional features, such as cilia length and beating frequency, together with targeted gene or marker analyses. Notably, we reanalyzed proximal and distal tissue datasets using published GEO raw or processed files, when available, to examine basal and secretory signatures. Representative profiles from one proximal (GSE150674) and one distal (GSE178360) dataset (Carraro et al. 2021; Kadur Lakshminarasimha Murthy et al. 2022) are shown in Figure S8A-B. In both datasets, clusters with canonical zone 1 secretory signatures (Carraro et al., “secretory 1”; Murthy et al., “MUC5AC⁺ MUC5B⁺”) showed the strongest correlation with secretory cells derived from proximal iBC-ALI and HBEC-ALI cultures. The presence of proximal-like secretory cells in the distal dataset reported by Murthy et al. (2022) is noted in the Table 1 footnote.

**Table 1.**
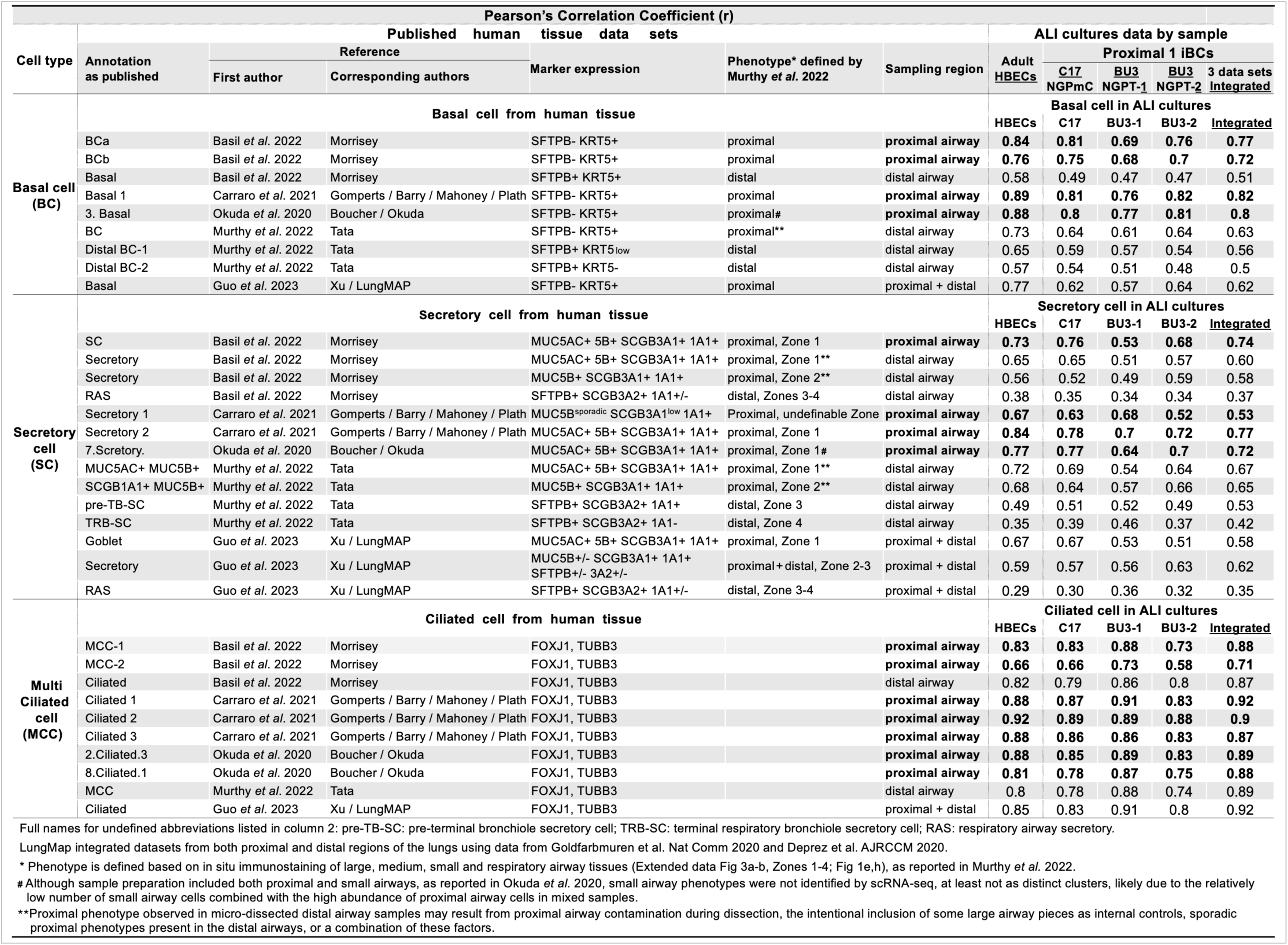
Proximal iBC-ALI and HBEC-ALI cultures contain basal and secretory cells that correlate closely with residential tissue basal and secretory cells in human proximal but not distal airways.

After examining the regional identity of major cell types in proximal 1 iBC and HBEC ALI cultures, we next assessed the maturation of proximal basal progenitors and iBCs at D45, D75, and D100 in 3D Matrigel droplets and at D130 in ALI cultures, alongside basal cell populations from HBEC ALI cultures, using published fetal lung/airway datasets (Miller et al. 2020; He et al. 2022; Hsu et al. 2024) and an adult proximal airway dataset (Carraro et al. 2021) as references. We also reanalyzed fetal tissue datasets to examine basal signatures. Representative profiles from a recent publication (He et al. 2022) and a recent bioRxiv preprint (Hsu et al. 2024) are shown in Figure S8C-D. Correlation analyses were performed using 4,500 HVGs per cell type across datasets. Correlations between three basal subpopulations from D45-130 and adult HBEC samples (Figures 2F and 4A-B, Basal progenitor 1/Basal 1/Basal; Basal progenitor 2/Basal 2/Cycling basal; Basal differentiating/Basal transitional) and corresponding adult proximal basal subpopulations (Figure S8A, Basal 1–3) increased progressively (Table 2). In contrast, correlations between these *in vitro* samples and fetal tissues were comparable across datasets (Miller et al. 2020; He et al. 2022; Hsu et al. 2024), with no significant differences observed (Table 2). Hsu et al. separately dissected tracheal, cartilaginous bronchi, non-cartilaginous airways, and distal airways from GW14.4–17.1 HFLs, and found that most basal subpopulations arose from proximal tracheal and cartilaginous bronchus regions, as shown in Figure 2C–F of the preprint (Hsu et al. 2024), consistent with our analysis (Figure S1A-B). Basal cells from all regions were combined in our correlation analysis. Notably, in He et al., “mid basal” and “late basal” cells were isolated from GW9–11 and GW15–22 samples, respectively, and these terms therefore reflect temporal rather than spatial differences (He et al. 2022). Together, these data indicate that region-specific analysis in currently available human fetal datasets is not feasible because non-cartilaginous and distal airway basal populations are extremely sparse. Our D45 samples did not preferentially align with fetal datasets, suggesting maturity exceeding that of GW5–22 fetal basal cells, or comparable maturity partially attenuated by Matrigel culture as a confounding factor. D75 and D100 samples demonstrated intermediate maturity, correlating more strongly with adult than fetal tissues. D130 iBC- and adult HBEC-ALI cultures exhibited the strongest alignment with adult references and exceeded correlations with all fetal datasets, indicating adult-like maturity.

**Table 2.** Progressive alignment of hPSC-derived basal subpopulations (D45–130) with adult proximal airway basal references, with comparable correlations across fetal tissues.

| Pearson's Correlation Coefficient (r) |  |  |  |  |  |  |  |  |  |  |  |  |  |  |  |  |  |
| --- | --- | --- | --- | --- | --- | --- | --- | --- | --- | --- | --- | --- | --- | --- | --- | --- | --- |
| Published human adult and fetal tissue data sets |  |  |  | hPSC-derived basal cells by day (D) of differentiation |  |  |  |  |  |  |  |  |  |  | Adult basal cell derived culture |  |  |
| Sample maturity | Sample region | Reference (First authors) (Last authors) | Annotation as published | D45 (UMAP in Fig 2F) |  | D75 (UMAP in Fig 2F) |  |  | D100 (UMAP in Fig 2F) |  |  | D130 (UMAP in Fig 4B) |  |  | HBECs (UMAP in Fig 4A) |  |  |
|  |  |  |  | Basal progenitor 1 | Basal progenitor 2 | Basal 1 | Basal 2 | Basal differentiating | Basal 1 | Basal 2 | Basal differentiating | Basal | Cycling basal | Basal transitional | Basal | Cycling basal | Basal transitional |
| Adult | Proximal | Carraro *et al.* 2021 Gomperts/Barry /Mahoney/Plath | Basal 1 | 0.58 | 0.36 | 0.63 | 0.35 | 0.22 | 0.66 | 0.37 | 0.34 | 0.82 | 0.41 | 0.49 | 0.89 | 0.46 | 0.58 |
|  |  |  | Basal 2 | 0.14 | 0.79 | 0.17 | 0.86 | -0.03 | 0.17 | 0.86 | -0.03 | 0.42 | 0.94 | 0.34 | 0.41 | 0.94 | 0.47 |
|  |  |  | Basal 3 | 0.51 | 0.31 | 0.35 | 0.39 | 0.47 | 0.45 | 0.33 | 0.46 | 0.49 | 0.45 | 0.83 | 0.53 | 0.51 | 0.86 |
|  |  |  | Basal progenitor 1 / Basal 1 / Basal correlates with Basal 1<br>Basal progenitor 2 / Basal 2 / Cycling basal correlates with Basal 2<br>Basal differentiating / Basal transitional correlates with Basal 3 |  |  |  |  |  |  |  |  |  |  |  |  |  |  |
| Fetal | Proximal + Distal | Hsu *et al.* 2024 Spence | LGR5+ basal 1 | 0.51 | 0.33 | 0.46 | 0.22 | 0.19 | 0.56 | 0.28 | 0.32 | 0.55 | 0.26 | 0.37 | 0.65 | 0.31 | 0.47 |
|  |  |  | LGR5+ basal 2 | 0.51 | 0.37 | 0.51 | 0.26 | 0.14 | 0.58 | 0.31 | 0.32 | 0.59 | 0.26 | 0.33 | 0.68 | 0.32 | 0.45 |
|  |  |  | Proliferating basal | 0.38 | 0.70 | 0.38 | 0.71 | -0.02 | 0.42 | 0.75 | 0.11 | 0.57 | 0.73 | 0.26 | 0.53 | 0.74 | 0.4 |
|  |  |  | Transitional basal | 0.52 | 0.29 | 0.38 | 0.21 | 0.3 | 0.52 | 0.25 | 0.4 | 0.44 | 0.27 | 0.55 | 0.53 | 0.35 | 0.65 |
|  |  |  | KRT13+ Epibasal | 0.42 | 0.16 | 0.19 | 0.13 | 0.39 | 0.34 | 0.13 | 0.38 | 0.21 | 0.21 | 0.68 | 0.27 | 0.3 | 0.7 |
|  |  |  | KRT4+ Epibasal | 0.13 | -0.03 | -0.15 | -0.01 | 0.52 | -0.02 | 0.07 | 0.25 | -0.08 | 0.05 | 0.54 | -0.03 | 0.12 | 0.48 |
| Basal progenitor 1 / Basal 1 / Basal correlates with LGR5+ basal 1 and 2<br>Basal progenitor 2 / Basal 2 / Cycling basal correlates with Proliferating basal<br>Basal differentiating / Basal transitional correlates with Transitional basal / KRT13+ and KRT4+ Epibasal |  |  |  |  |  |  |  |  |  |  |  |  |  |  |  |  |  |
| Fetal | Proximal + Distal | He *et al.* 2022 | Proximal basal | 0.51 | 0.31 | 0.42 | 0.27 | 0.16 | 0.53 | 0.26 | 0.37 | 0.5 | 0.28 | 0.42 | 0.6 | 0.36 | 0.53 |
|  |  |  | Mid basal | 0.55 | 0.44 | 0.52 | 0.36 | 0.08 | 0.61 | 0.39 | 0.30 | 0.6 | 0.37 | 0.39 | 0.62 | 0.4 | 0.46 |
|  |  |  | Late basal | 0.44 | 0.36 | 0.44 | 0.34 | 0.09 | 0.48 | 0.35 | 0.29 | 0.52 | 0.34 | 0.34 | 0.57 | 0.38 | 0.45 |
| Basal progenitor 1 / Basal 1 / Basal from each sample show similar Correlation with all 3 populations from this fetal data set. |  |  |  |  |  |  |  |  |  |  |  |  |  |  |  |  |  |
| Fetal | Proximal + Distal | Miller *et al.* 2020 | Basal | 0.52 | 0.36 | 0.34 | 0.36 | 0.32 | 0.50 | 0.36 | 0.46 | 0.41 | 0.44 | 0.67 | 0.52 | 0.52 | 0.71 |
| Basal progenitor 1 / Basal 1 / Basal, Basal progenitor 2 / Basal 2 / Cycling basal from each sample show similar Correlation with Basal.<br>Basal transitional from D130– and adult HBEC–derived cultures correlate better with Basal. |  |  |  |  |  |  |  |  |  |  |  |  |  |  |  |  |  |

**Correlation with adult proximal basal populations increased progressively from D45 to D130:**
- o D130 iBC-ALI and adult HBEC-ALI cultures showed the strongest alignment with adult references and exceeded correlations with all fetal datasets, indicating adult-like maturity. - o D75 and D100 samples demonstrated intermediate maturity, correlating more strongly with adult than fetal tissues. - o D45 samples showed comparable correlations with adult and fetal datasets.
**Overall, in vitro samples showed comparable correlations across fetal tissues:**
- o D45 did not preferentially align with fetal datasets, suggesting maturity exceeding that of GW13–23 fetal basal cells, or comparable maturity partially attenuated by Matrigel culture as a confounding factor. - o D130 and adult HBEC ALI samples did not exhibit reduced correlations with fetal tissues relative to D45–D100 samples, consistent with fewer culture-specific artifacts under ALI conditions compared with 3D Matrigel culture.
**Note:** In Hsu *et al.*, most basal subpopulations arose from proximal tracheal and cartilaginous bronchial regions, and basal cells from all regions were combined in the present correlation analysis.

Lastly, because ionocytes are proximally enriched across species (Okuda et al. 2020; Travaglini et al. 2020; Yuan et al. 2023), we further evaluated ionocytes in iBC-derived ALI cultures across conditions at transcriptomic and protein levels to verify preferential ionocyte specification in proximal conditions. First, we reanalyzed ionocyte signatures in individual scRNA-seq datasets across all conditions using an independent pipeline developed by the Cystic Fibrosis Foundation Therapeutics (CFFT) lab. Specifically, we evaluated primary HBEC-derived ALI cultures from seven subjects (Carraro et al. 2021), together with ALI cultures derived from region-specific iBCs and CSC protocol iBCs. Ionocyte signatures were defined as the top 50 DEGs identified by scRNA-seq in HBEC ALI cultures in the original study (Plasschaert et al. 2018). Ionocyte signatures were readily detected in primary HBEC and region-specific iBC samples but were absent from all seven iBC-CSC protocol samples (Figure 6A). Notably, this pipeline also identified the few ionocytes in intermediate iBC-derived ALI culture (Figure 6A) that clustered with basal transitional cells in Figures 4C and S7C. Across both scRNA-seq analyses, ionocyte signatures were consistently detected in HBEC- and proximal iBC-derived cultures (Figures 4A–C and 6A), yet ionocytes were markedly underrepresented by scRNA-seq. In a recent ferret genetic model, ionocyte damage during 10X Genomics capture was quantified. With ∼10,000 live FOXI1–eGFP⁺ as input cells, only 434 ionocytes were identified (Yuan et al. 2023). Accordingly, we quantified ionocytes in each region-specific iBC-derived ALI culture by IF for FOXI1 protein. Ionocytes were consistently reduced in intermediate iBC-ALI cultures relative to proximal conditions (paired *t* test, P < 0.01 and P < 0.005), whereas the distal condition showed rare but detectable ionocytes (Figure 6B). These findings aligned with prior reports that ionocytes are enriched in proximal large airways and sparse in distal small airways of adult human lungs (Okuda et al. 2020; Travaglini et al. 2020; Yuan et al. 2023). In parallel CSC-protocol iBC-derived ALI cultures, sporadic FOXI1⁺ cells were identified (Figure 6B). Because the CSC protocol relies on endogenous BMP and WNT signaling, with CHIR withdrawn from D15 onward, it would be expected to yield iBCs resembling intermediate airway populations and therefore partially overlapping phenotypically with our intermediate iBC condition.

**Figure 6.**
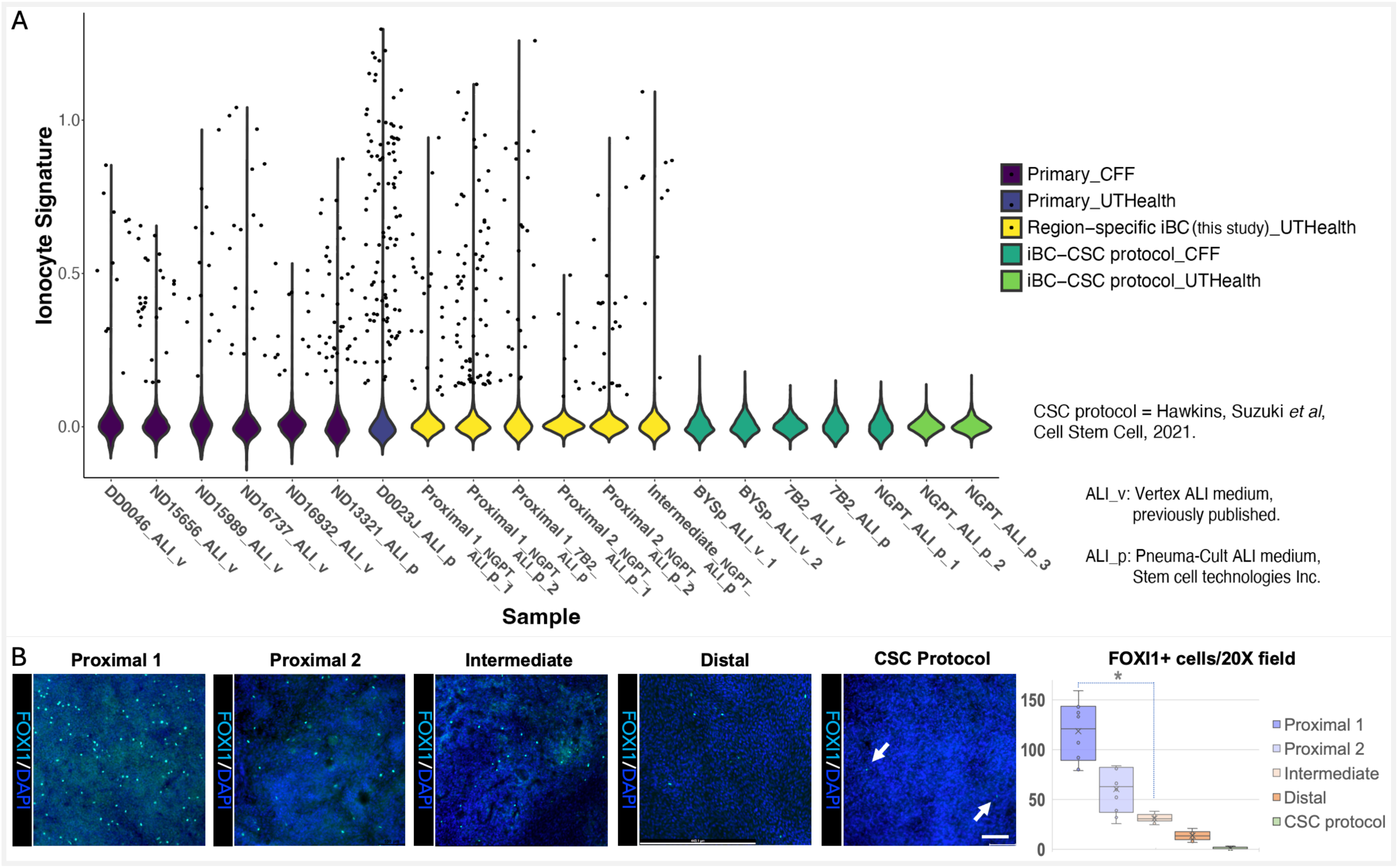
Characterization of ionocytes in region-specific iBC-ALI cultures. (**A**) Signatures of cells classified as ionocytes in the indicated ALI culture samples. CFF indicates that both the experiment and scRNA-seq were performed in the Cystic Fibrosis Foundation Therapeutics (CFFT) lab, whereas UTHealth indicates that both were performed at UTHealth. UMAPs of the integrated and/or individual datasets for each condition are shown in Figures 4 B–D and 4E. Analysis was performed using a pipeline established at the CFFT lab, with the top 50 DEGs used as input. Primary HBEC samples are labeled by sample ID followed by the ALI medium used. iBC-CSC samples are labeled by cell line_ALI medium_biological replicate number. Region-specific iBC samples are labeled by condition_cell line_ALI medium_biological replicate number. Primary HBECs and region-specific iBC samples were differentiated in ALI medium for 21-30 days. iBC-CSC protocol samples were differentiated in ALI medium for ∼14 days, as cultures generated under this condition tended to collapse when maintained for longer than 14 days. (**B**) D90+ region-specific iBCs and iBCs generated using the CSC protocol were differentiated in ALI medium for 30 and 14 days, respectively, and stained for FOXI1. *n* = 3 for each hPSC line; a representative experiment from BU3 NGPT is shown. The number of FOXI1^+^ cells per 20× microscopic field was quantified from two BU3 NGPT experiments and plotted as shown. Arrows indicate rare FOXI1^+^ cells. Scale: 100μm.

### Proximal iBCs derived from G551D *CFTR* hiPSCs recapitulate the CF ionocyte phenotype

Studies have shown that ionocytes and secretory cells are the main CFTR-expressing populations, with ionocytes, although rare, exhibiting the highest per-cell CFTR expression (Montoro et al. 2018; Okuda et al. 2020; Plasschaert et al. 2018; Travaglini et al. 2020; Yuan et al. 2023), consistent with our findings in HBEC- and proximal iBC-ALI cultures (Figure 3A–B). In a ferret genetic model (Yuan et al. 2023), ionocytes contribute ∼50% of proximal airway CFTR activity and are required for normal physiology. In addition, cultured airway samples from patients with CF exhibit increased ionocyte frequency due to a feedback compensation mechanism mediated by SHH signaling (Cai et al. 2023). Given the robust ionocyte potential of proximal iBCs, we leveraged this system to recapitulate this CF-associated phenotype using G551D *CFTR* hiPSCs. G551D is a Class III mutation that impairs CFTR channel gating and reduces open probability (Welsh and Smith 1993; Rowe, Miller, and Sorscher 2005; Veit et al. 2016). We used gene-edited G551D C17 NGPmC hiPSCs, which are isogenic to the C17 NGPmC dual-reporter line, harbor the Gly551Asp variant (c.1652G>A, p.Gly551Asp), and carry the NKX2-1^GFP^ TP63^mCherry^ reporters (gift from John Mahoney, Cystic Fibrosis Foundation, and previously published) (Berical et al. 2022). Proximal 1 and 2 iBCs derived from isogenic C17 hiPSCs carrying either wild type (WT) or G551D *CFTR* were differentiated at ALI for 4–5 weeks. G551D C17-derived proximal 1 and 2 cultures contained, on average, 2.65 ± 0.64 to 2.88 ± 0.82 fold more FOXI1+ cells per 20x field, respectively, than WT C17-derived ALI cultures (P < 0.005) (Figure 7A–C). The BSND^+^ fraction among FOXI1^+^ cells ranged from ∼44–67% across conditions (Figure 7C), with no significant difference between G551D and WT lines. Notably, under WT Proximal 1 and 2 conditions, the C17 line exhibited fewer FOXI1⁺ cells than BU3 (Figure 6B), suggesting that line-specific differences influence ionocyte output, potentially including higher endogenous BMP signaling in C17 than in BU3. Together, these data demonstrate that proximal iBCs can recapitulate the ionocyte phenotype observed in CF airways. Future studies will extend this platform to recapitulate mechanisms underlying additional CF-related phenotypes, including the relative contribution of ionocytes to CFTR activity in humans, as well as a broader range of genetic and non-genetic airway diseases.

**Figure 7.**
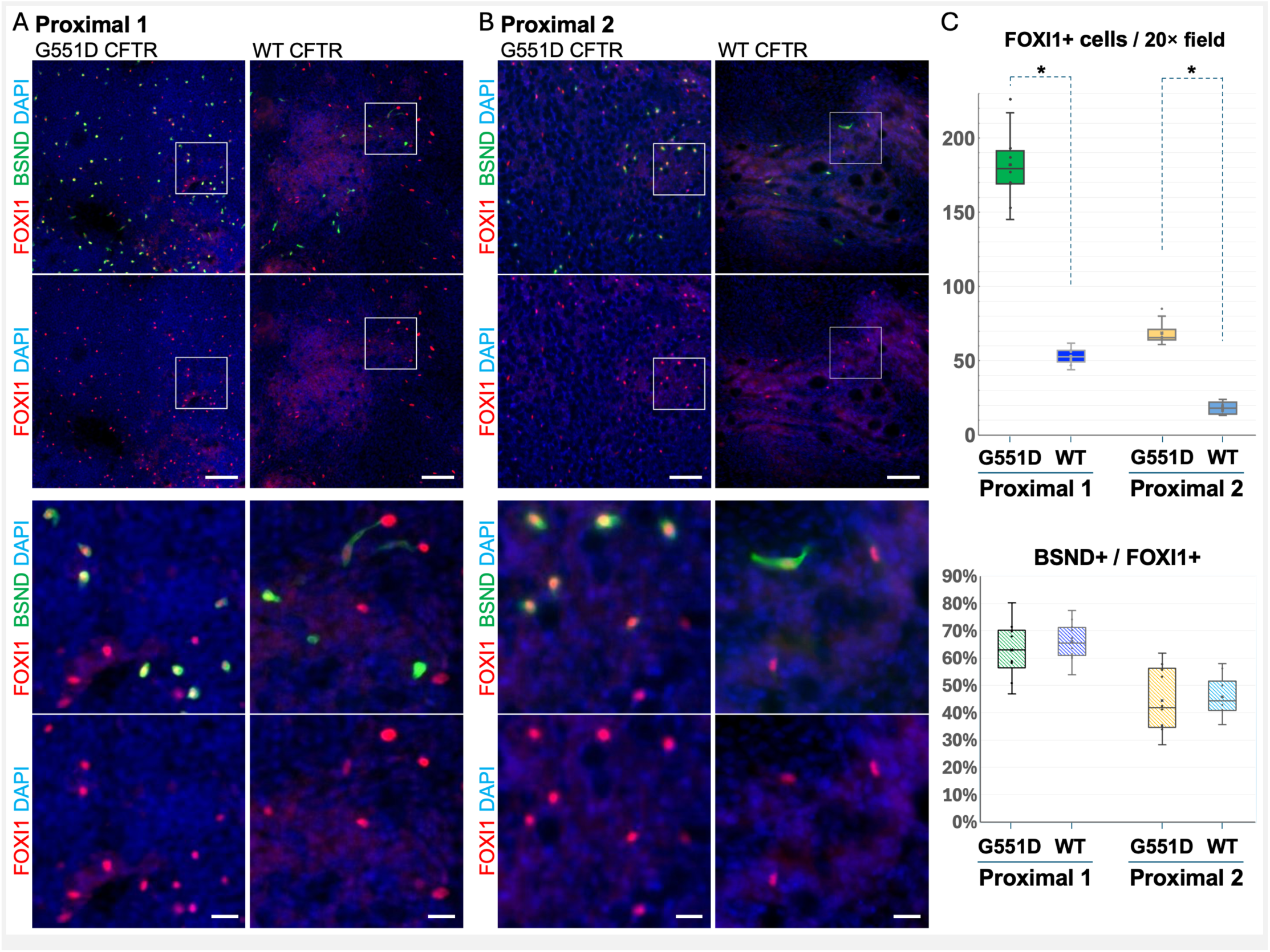
Characterization of ionocytes in proximal iBC–ALI cultures derived from isogenic WT and G551D *CFTR* C17 NGPmC hiPSCs. (**A-B**) D90+ proximal 1 and proximal 2 iBCs derived from isogenic C17 hiPSCs carrying WT or G551D *CFTR* mutation were differentiated in ALI medium for 4–5 weeks and stained for FOXI1 and BSND (*n* = 3 per line). Insets for each condition are shown in the bottom panels. Scale bars (*top*): 100 µm; insets (*bottom*): 20 µm. (**C**) FOXI1⁺ and BSND^+^/FOXI1^+^ cell populations per 20× field were quantified from 10 random fields across two experiments and plotted as shown. The two indicated bracketed comparisons were analyzed independently; *P<0.005. Notably, the frequency of FOXI1⁺ cells under proximal 1 and 2 conditions in the WT C17 NGPmC line was lower than in BU3 NGPT (Figure 6B), potentially reflecting higher endogenous BMP production in C17.

## Discussion

hPSC-derived lung models, including mixed-lineage organoids and purified specialized cell types, provide valuable platforms for recapitulating human lung development, modeling disease, and advancing regenerative medicine. Both approaches begin by deriving early lung progenitors that resemble primordial bud-tip progenitors capable of generating any airway and alveolar lineages. Recent studies report significant progress in generating mixed-lineage cultures (Mou et al. 2012; Huang et al. 2014; Firth et al. 2014; Gotoh et al. 2014; Dye et al. 2015; Rankin et al. 2016; Konishi et al. 2016; Chen et al. 2017; Hawkins et al. 2017; McCauley et al. 2017; Jacob et al. 2017; Yamamoto et al. 2017; Miller et al. 2018; Miao et al. 2025; Vila-Gonzalez et al. 2024). In contrast, generating purified specialized cell types requires more precise recapitulation of lineage-specific developmental pathways. Although challenging, this enables rigorous phenotypic and functional characterization essential for downstream applications, particularly cell-based therapies. Motivated by this goal and recent progress alongside unmet challenges (Hawkins et al. 2021; Wang et al. 2023), we generated regionally defined airway basal cells – the principal stem cells of the conducting airways.

Here we demonstrate directed differentiation of hiPSCs into region-specific iBCs that closely resemble the molecular and functional phenotypes of adult human proximal, intermediate, and distal airway BCs. Notably, the generation of proximal iBCs has not previously been achieved. We show that proximal iBC–derived ALI cultures exhibit CFTR anion-channel activity comparable to that of HBEC–derived cultures. We further demonstrate that proximal iBCs generate the full repertoire of airway cell types, including ionocytes, *in vitro* and *in vivo* in a tracheal xenograft model, as well as their value for modeling human airway development and disease.

Our approach overcomes key constraints in hPSC airway differentiation, including maintenance of airway-progenitor lung fate and the regional identity, maturity, and differentiation competence of iBCs (Firth et al. 2014; Konishi et al. 2016; McCauley et al. 2017; Hawkins et al. 2021; Wang et al. 2023). We achieved this by properly recapitulating essential developmental cues. Drawing on our prior work in hPSC differentiation, we previously showed that lung-progenitor specification from hPSCs requires three indispensable pathways: WNT, BMP, and RA (Huang et al. 2014). We therefore reasoned that downstream airway basal-cell specification similarly relies on combined inputs from essential developmental pathways rather than a single cue. During development, proximal–distal patterning coincides with airway basal specification; we infer that these processes are governed by distinct yet coordinated pathways. Prior work and the CSC protocol primarily modulated WNT signaling, showing that WNT withdrawal is essential for basal specification but contributes little to proximal–distal regional patterning (Hawkins et al. 2021; McCauley et al. 2017). However, existing studies neither explicitly modulated proximal–distal patterning after lung progenitor specification nor systematically characterized the regional identity of the resulting airway lineages. Accordingly, our study centered on regional patterning while also addressing lung-fate maintenance, maturation, and differentiation competence – limitations observed with CSC-derived iBCs. Using our proximal–distal patterning roadmap as a readout, we specified proximal-distal regional identity by BMP inhibition with NOGGIN or BMP activation with BMP4. Cultures defaulted toward an intermediate identity under endogenous BMP signaling. Our data support the concept that the NOGGIN–BMP axis is a principal regulator of proximal–distal patterning in hiPSC-derived lung progenitors. This finding aligns with mouse *in vivo* studies (Lu et al. 2001; Weaver et al. 1999), and may reflect conserved roles of BMP signaling in proximal–distal airway patterning in humans and mice, while allowing for species-specific differences in cellular composition and anatomical structure. In parallel, we applied temporally tapered WNT activation to balance the seemingly opposing requirements for reduced WNT during basal specification and sustained WNT activity for lung-fate maintenance, thereby recapitulating the critical role of signaling timing in endodermal development *in vivo* (Rankin et al. 2018). Together, coordinated NOGGIN–BMP modulation and tapered WNT signaling efficiently establish regional identity while sustaining lung fate during airway basal specification. More broadly, our findings suggest that BMP and WNT pathways, which are required for early lung progenitor specification (Rankin et al. 2016; Huang et al. 2014; Hawkins et al. 2017; Chen et al. 2017), can be redeployed at later stages of hPSC differentiation to achieve stage-specific outcomes, consistent with mouse lung development *in vivo* (Que et al. 2006; Li et al. 2008; Goss et al. 2009; Morrisey and Hogan 2010; Weaver et al. 1999; Lu et al. 2001; Mucenski et al. 2003; Shu et al. 2005; Hashimoto et al. 2012; Hou et al. 2019).

Beyond lung fate maintenance, WNT activation during D15–45 likely promoted iBC maturation and differentiation competence. Comparison of our intermediate iBC condition with the CSC protocol supports the conclusion that both regimens yield iBCs with an intermediate regional identity driven by endogenous BMP signaling, but CSC-derived iBCs require commercial medium to induce the mature basal marker NGFR, express low levels of KRT5, and generate secretory and ciliated progeny resembling human fetal lung counterparts, with variable ionocyte differentiation and absent PNECs in iBC-ALI cultures (Hawkins et al. 2021; Wang et al. 2023). The ionocyte output reported by Wang et al. may reflect repeated NGFR-based enrichment of the subset of iBCs competent for ionocyte differentiation, rather than efficient proximal patterning of the culture as a whole. Because our intermediate iBC and the CSC protocol use the same strategy to generate NKX2-1^+^ lung progenitors from hPSCs through WNT, BMP, and RA activation (Huang et al. 2014; Hawkins et al. 2021) and they differ primarily in the timing of subsequent WNT withdrawal. The additional WNT exposure during the D15–45 window likely contributes to the improved iBC maturation and competence of our intermediate iBCs. To focus specifically on NOGGIN–BMP–mediated proximal–distal patterning, we otherwise matched the CSC protocol baseline conditions, reducing only FGF10, cAMP, and IBMX to minimize potential confounders. Although higher FGF10 can enhance basal proliferation in hPSC differentiation culture (Hawkins et al. 2021), its potential to promote distalization through endogenous BMP signaling should be considered, given its role in branching morphogenesis and distal lung development (Hyatt, Shangguan, and Shannon 2004; Weaver, Dunn, and Hogan 2000; Bellusci et al. 1997; Yuan et al. 2018). Likewise, because glucocorticoid signaling promotes distal lung maturation and alveolar gene expression (Bird et al. 2015; Cole et al. 2004), we reduced cAMP and IBMX to avoid potential distalization. Although these additional factors may also contribute, they are less likely than WNT timing to account for the observed differences, a possibility that will require further study.

As proof of principle, disease modeling with CSC protocol–derived iBCs has previously demonstrated IL-13–induced mucus metaplasia, F508del *CFTR* mutation–induced chloride-channel dysfunction with partial restoration, and DNAH5 mutation-associated ciliary dysfunction (Hawkins et al. 2021). Here, leveraging the robust ionocyte potential of proximal iBCs, we recapitulated the ionocyte phenotype observed in CF airways (Cai et al. 2023) using G551D *CFTR* hiPSCs. This platform is therefore well positioned to dissect additional CF-related phenotypes and mechanisms, particularly the debated relative contributions of secretory cells versus ionocytes to CFTR function (Okuda et al. 2020; Yuan et al. 2023). Our region-specific iBCs recapitulate regional secretory cell phenotypes (Naizhen et al. 2019; He et al. 2022; Kadur Lakshminarasimha Murthy et al. 2022) and match proximal adult BCs in ionocyte potential (Figure 3I–J), providing a unique opportunity to address this question. More broadly, the ability to generate rare cell types creates opportunities to study diverse genetic and non-genetic airway diseases and to probe rare-cell biology. Notably, tuft cells, although proximally enriched *in vivo* (Montoro et al. 2018; Plasschaert et al. 2018), were detected only sporadically by scRNA-seq and did not form a distinct cluster in either adult HBEC-derived or proximal iBC-derived cultures, suggesting that standard ALI media may be suboptimal for tuft differentiation. Consistent with this, a recent study (Shah et al. 2025) showed that tuft cells are rare both *in vivo* at steady state and in ALI cultures, but increase upon specific cytokine stimulation. These possibilities will be examined in future studies.

### Limitations of the Study

Lineage-tracing studies in the developing mouse lung show that only a subset of TP63⁺ cells commits to a basal fate postnatally, whereas the remaining traced cells downregulate TP63 and adopt secretory or ciliated fates (Yang et al. 2018). Consistent with this, under intermediate and distal conditions, a fraction of initially sorted TP63⁺ cells downregulated TP63 and assumed a differentiating airway progenitor phenotype, with TP63 fate stabilizing around D75–90. By contrast, under proximal conditions with NOGGIN plus tapered WNT, most proximal TP63⁺ airway progenitors in our system preferentially established a basal fate. This phenomenon aligns with the decreasing abundance of basal cells from proximal to distal airways. Given the robust generation of proximal iBCs, the present study focuses on in-depth characterization of this population, which, to our knowledge, has not previously been generated. Conversely, the generation of hPSC-derived intermediate and distal iBCs is less efficient, although these cells can still be produced in large numbers *in vitro*, potentially facilitating studies of regional BCs given the relative scarcity and limited availability of basal cell populations in developing intermediate and distal airways in human fetal lungs (Hsu et al. 2024). Nevertheless, efficient intermediate and distal basal specification, particularly under distal conditions, likely requires additional signaling inputs beyond BMP and WNT. Future work will be needed to define these inputs, including whether NOGGIN actively promotes and maintains basal identity over alternative airway lineages.

Under high-level WNT activation, adding NOGGIN or BMP4 during D15–D30 reduced maintenance of WNT-driven NKX2-1 in a subset of cells, indicating cross-talk between WNT and the NOGGIN–BMP axis in NKX2-1 regulation. Prior work suggests BMP does not directly regulate the *NKX2-1* promoter (Domyan et al. 2011), indicating an indirect mechanism. Our data show that NOGGIN and BMP4 directly upregulate *SOX2* and *SOX9*, respectively, and that WNT effector TCF4 binds the *SOX2* locus. We also previously showed that SOX9 expression in lung progenitors is WNT-dependent (Ostrin et al. 2018). Together, these observations suggest that elevated SOX2 and SOX9 may indirectly modulate NKX2-1 under proximal and distal conditions, respectively; future studies will define the mechanism.

While the tracheal xenograft model effectively assesses the *in vivo* differentiation capacity of iBCs, it does not fully recapitulate the airway reconstitution recently achieved by transplantation in immunodeficient mice (Ma et al. 2023), a step closer to clinical translation. That study evaluated CSC protocol–derived iBCs (from mESCs and hiPSCs). Encouragingly, our iBCs exhibit greater maturation and competence than CSC-derived cells. Evaluating region-specific iBCs in this transplantation system will be an important next step toward assessing their potential for cell-based therapy.

## Experimental Materials and Methods

### Human subjects

De-identified human bronchial epithelial cells (HBECs) were generously provided by Dr. Scott Randell and the CF Center Tissue Procurement and Cell Culture Core at the University of North Carolina. De-identified human fetal lung (HFL) tissue was obtained under IRB protocol No. STUDY-22-00065, approved by the Icahn School of Medicine at Mount Sinai, and collected by the Developmental Origins of Health and Disease Biorepository (DOHaD) at Mount Sinai. Tissue samples were processed for frozen sections by Dr. Ya-Wen Chen’s group at Mount Sinai. Both HBECs and HFL tissue samples are exempt from regulation by the Committee for the Protection of Human Subjects at the University of Texas Health Science Center at Houston (UTHealth) under 45 CFR 46.101(b).

### Animals

Adult mice (Foxn1^nu^; Jackson Laboratory) and rats (NIH-Foxn1^rnu^; Charles River) were housed under specific pathogen–free conditions in the AAALAC-accredited barrier facility of the Brown Institute of Molecular Medicine at UTHealth. Animal rooms were maintained at 21–23 °C on a 12:12-h light:dark cycle. Animals were housed in autoclaved, individually ventilated cages with irradiated corncob bedding and cotton square nestlets and were provided an irradiated, balanced diet ad libitum (5058; LabDiet, St. Louis, MO) and acidified water. Animals were group-housed before procedures; mice receiving xenografts were singly housed after surgery. All procedures were approved by the UTHealth IACUC and conducted in accordance with the NIH Guide for the Care and Use of Laboratory Animals.

### Primary HBEC maintenance

Cryopreserved HBECs were thawed and expanded on 804G-conditioned medium-coated tissue-culture plates (Mou et al. 2016) in PneumaCult-Ex Plus (STEMCELL Technologies, Inc., Vancouver, Canada) supplemented with 1 μM A83-01, 1 μM DMH1, and 10 μM Y-27632 (all from Tocris Bioscience, Bio-Techne Corporation, Minneapolis, MN). At 60–70% confluence, HBECs were dissociated with Accutase (Innovative Cell Technologies, Inc., San Diego, CA) and seeded onto 804G-conditioned medium-coated 6.5 mm polyester membrane Transwells inserts with 0.4 μm pore size (Corning Incorporated, Corning, NY) at 100,000–150,000 cells per insert, maintained submerged in the same medium for 48 h before initiating air–liquid interface (ALI) differentiation.

### hPSC Maintenance

hESC line RUES2 (Rockefeller University Embryonic Stem Cell Line 2; NIH registration number 0013; a generous gift from Ali Brivanlou, Rockefeller University), RUES2 NKX2-1^GFP^, and RUES2 NKX2-1^GFP^ SOX2^mRUBY3^ were maintained on irradiated mouse embryonic fibroblasts (∼25,000 cells/cm^2^, Gibco, Thermo Fisher Scientific Inc., Waltham, MA) in DMEM/F12 (Corning), 20% knockout serum replacement (Gibco), 0.1 mM β-mercaptoethanol (MilliporeSigma, St. Louis, MO) and 20 ng/ml FGF-2 (R&D Systems, Bio-Techne Corporation, Minneapolis, MN) as previously described (Huang et al. 2015; Huang et al. 2014). All hiPSC lines were maintained in feeder-free conditions on hESC-qualified Matrigel (Corning) coated 6-well plates (Corning) in StemFlex Medium (Thermo Fisher Scientific) or mTeSR1 medium (STEMCELL Technologies). Cultures were maintained in an undifferentiated state in a 5% CO_2_/air environment. Both hESC and hiPSC lines are passaged with Accutase or ReLeSR (STEMCELL Technologies). All hESC and hiPSC lines used were characterized for pluripotency, found to be karyotypically normal; and routinely checked for mycoplasma contamination.

### Dual fluorescent reporter hESC and hiPSCs

The dual reporter RUES2 line expressing NKX2-1^GFP^ and SOX2^mRUBY3^ (R2 NGSmR) was derived from RUES2 hPSCs using CRISPR/Cas9. NKX2-1-GFP construct was generated at UTHealth and previously published (Hawkins et al. 2017). A gRNA targeting exon 1 of endogenous *SOX2* was designed against the DNA target sequence 5′-CTGCCCCCTCTCACACATGG-3′, and a donor template containing the mRUBY3 sequence was constructed. In this report, we used the clones R2 NGSmR c3 and c5. Additional dual reporter lines, including NKX2-1^GFP^ TP63^tdTomato^ BU3 hiPSCs (BU3 NGPT) (Hawkins et al. 2021), NKX2-1^GFP^ TP63^mCherry^ C17 hiPSCs with and without G551D *CFTR* mutation (C17 NGPmC, G551D C17 NGPmC; a generous gift from John Mahoney, Cystic Fibrosis Foundation) (Berical et al. 2022) were previously described.

### Differentiation of hPSCs into lung progenitors

NKX2-1^+^ lung progenitors were generated through the stepwise differentiation of hPSCs into definitive endoderm, anterior foregut endoderm, and lung progenitors, using previously published approaches with modifications. Previously, before definitive endoderm induction, we treated the undifferentiated hPSCs with a low dose of human BMP4 (3 ng/ml) for 24 hr to mimic the primitive streak formation in embryonic development. We were able to skip this step without affecting the differentiation efficiency for all the cell lines used in the current study. For most of the experiments, definitive endoderm induction was performed in serum-free differentiation (SFD) medium of DMEM/F12 (3:1) (Corning) supplemented with N2 Supplement (Thermo Fisher), B27 Supplement with retinoic acid (Thermo Fisher), ascorbic acid (50 μg/ml, MilliporeSigma), Glutamax (2 mM, Thermo Fisher), monothioglycerol (MilliporeSigma), 0.05% bovine serum albumin (BSA) (Thermo Fisher), 1% penicillin-streptomycin (Corning). hPSCs were briefly dissociated into 2–10 small cell clusters and plated onto low attachment 6-well plates to form embryoid bodies in SFD media containing Y-27632, 10 μM, human BMP4, 0.5 ng/ml (R&D Systems); human bFGF, 2.5 ng/ml; human Activin A, 100 ng/ml (R&D Systems) for 64–72h. Cells were fed every 36–48 h (depending on the density) by removing half of the old media and adding half fresh media. For some experiments in BU3 NGPT line, definitive endoderm induction was achieved with culture in STEMdiff Definitive Endoderm Kit (STEMCELL Technologies) for 64–72 hours. For anterior foregut endoderm induction, endodermal cells were dissociated into single cells/small clusters using accutase (3–4 min) and plated on Matrigel-coated 6 or 12-well tissue culture plates (∼50,000 cells/cm^2^) in SFD medium supplemented with 100 ng/ml human NOGGIN (R&D Systems) and 10 μM SB431542 (Tocris) for 24 h, and then switched to 2 μM Dorsomorphin dihydrochloride (Tocris), 10 μM SB431542 for 40–48 h. NOGGIN is a biological inhibitor of BMP, while Dorsomorphin dihydrochloride is a pharmacological inhibitor of BMP. The purpose of using NOGGIN for the first 24 hours is to avoid Dorsomorphin-related cellular toxicity, as the cells are more sensitive due to stress and toxicity induced by enzymatic dissociation. The total duration for anterior foregut endoderm induction is 64–72 h. For some hiPSC lines, a low dose of Y-27632 (2.5 μM) was added along with 100 ng/ml human NOGGIN and 10 μM SB431542 during the first 24 hours of anterior foregut endoderm induction to mitigate stress and toxicity induced by enzymatic dissociation. For lung progenitor induction, the resulting anterior foregut endoderm was treated with a ventralization cocktail containing CHIR99021, 3 μM (WNT signaling agonist, Tocris); human FGF10, 10 ng/ml (R&D Systems); human BMP4, 10 ng/ml and all-trans retinoic acid (ATRA), 50–100 nM (MilliporeSigma) in SFD media for 9–10 days. At D15, NKX2-1^+^ lung progenitors were sorted based on NKX2-1^GFP^ expression for reporter lines BU3 NGPT, C17 NGPmC, and RUES2 NGSmR, or expression of the cell surface marker CPM (anti-CPM, Cat.# 014-27501, FUJIFILM Wako Pure Chemical Corporation) and EpCAM (anti-EpCAM, Alexa Fluor 647-conjugated, Cat.# 324212, BioLegend Inc., San Diego, CA) for non-reporter hPSC lines.

### Induction of airway progenitors from hPSC-derived lung progenitors

Sorted D15 NKX2-1^+^ or CPM^+^ EpCAM^+^ lung progenitors were embedded in growth factor reduced Matrigel (Corning) for three-dimensional (3D) culture at a density of 400 cells per μl, in 40–50 μL droplets. Each droplet was placed onto the surface of a warm tissue-culture plate and incubated at 37 °C for 20–30 min to allow polymerization. The 3D culture was carried out in SFD medium containing: (1-2) 3 μM CHIR99021, 10 ng/ml FGF10, low, medium or high Noggin and 5 μM Y-27632 for 15 days to induce proximal airway progenitor; (3) 3 μM CHIR99021, 10 ng/ml FGF10 and 5 μM Y-27632 for 15 days for intermediate multipotent progenitor induction; (4) 3 μM CHIR99021, 10 ng/ml FGF10, low BMP4 and 5 μM Y-27632 for 30 days for distal multipotent progenitor induction. Distal cells were analyzed and re-embedded at both D30 and D45. Three additional conditions were used side-by-side as controls: (5) 250 ng/ml <u>FGF2</u>, 100 ng/ml FGF<u>10</u>, 50nM <u>d</u>examethasone, 100 μM 8-Bromoadenosine 3′,5′-<u>c</u>yclic monophosphate sodium salt (<u>c</u>AMP; Tocris), 100 μM 3-Isobutyl-1-methylxanthine (IBMX; Tocris), and 10 μM <u>Y</u>-27632, abbreviated as FGF2+10+DCI+Y – this is our previously published (Hawkins et al. 2021) airway basal condition. FGF2 promotes cell growth, while DCI has been used to mature lung epithelial cells by us and others; (6) condition 5 + NOGGIN; (7) condition 5 + BMP4. Notably, the main purpose of conditions 5–7 was to serve as no-WNT-activation (minus CHIR) controls for each of our experimental conditions (1–4). Compared to plain no-WNT-activation controls, conditions 5–7 had a higher dosage of FGF10 and included FGF2 and DCI, all of which significantly promote cell proliferation. In contrast, without CHIR, lung progenitors in plain no-WNT-activation control conditions (FGF10+NOGGIN+Y, FGF10+Y, and FGF10+BMP4+Y) exhibit minimal growth. Despite differences in cell growth, our tests indicate that conditions 5–7 and the plain no-WNT-activation controls display similar lung fate maintenance phenotypes. At D30/45, airway and multiplepotent progenitors and the cells from control conditions were analyzed for proximal-distal patterning by NKX2-1, SOX2, SOX21 and SOX9 expression, and sorted based on NKX2-1^GFP^ or CPM and EpCAM expression for further culture.

### Differentiation of airway progenitors into basal progenitors

Sorted D30/45 airway and multipotent progenitors were embedded in growth factor reduced Matrigel for 3D culture in SFD medium containing (1-2) 1 μM CHIR99021, 10 ng/ml FGF10, low, medium or high NOGGIN and 10 μM Y-27632 for 15 days to induce proximal basal progenitors; (3) 1 μM CHIR99021 and 10 μM Y-27632 for 15 days for intermediate basal progenitor induction; (4) 1 μM CHIR99021, 10 ng/ml FGF10, low BMP4 and 10 μM Y-27632 for 15 days for distal basal progenitor induction. Sorted NKX2-1^+^ cells from the three control conditions were carried further in SFD medium containing: (5) FGF2+10+DCI+Y – our previously published airway basal condition; (6) condition 5 + NOGGIN; (7) condition 5 + BMP4. At D45/60, basal progenitors and the cells from control conditions were analyzed for basal specification efficiency and sorted based on NKX2-1^GFP^ p63^tdTomato^, NKX2-1^GFP^ TP63^mcherry^ reporter expression or enriched based on surface markers F3 (anti-F3, Cat.# 365204, BioLegend), EGFR (anti-EGFR, Cat.# 352907, BioLegend) and PDPN (anti-PDPN, Cat.# 337024, BioLegend) co-expression (for non-reporter hPSC lines) for further culture.

### Basal progenitor maturation and basal cell expansion and maintenance

Sorted D45/60 basal progenitors were embedded in growth factor reduced Matrigel for 3D culture in SFD medium containing: (1-2) 10 ng/ml FGF10, low, medium, or high NOGGIN, 250 ng/ml FGF2, 50nM dexamethasone, 100 μM cAMP, 100 μM IBMX and 10 μM Y-27632 for proximal basal maturation, expansion, and maintenance; (3-4) 10 ng/ml FGF10, 250 ng/ml FGF2, 50nM dexamethasone, 100 μM cAMP, 100 μM IBMX and 10 μM Y-27632 for intermediate and distal basal maturation, expansion, and maintenance. As explained in Results section, in proximal condition, NOGGIN is required during maturation due to the continuous production of endogenous BMPs detected in basal progenitors. In distal condition, BMP4 is withdrawn to avoid excessive suppression on SOX2. In some experiments with BU3 NGPT line, the cells were cultured in basal progenitor induction medium for additional 15 days and switched to basal maturation medium at D60/75. Cells from the control conditions were cultured in SFD medium containing: (5a) FGF2+10+DCI+Y–these medium components cannot mature the basal cells generated using this method; (5b) PneumaCult ExPlus supplemented with 1 μM A83-01, 1 μM DMH1, and 10 μM Y-27632, referred to as ’Basal Cell Medium’ for CSC protocol (Hawkins et al. 2021) and used to mature iBCs generated using this method; (6) condition 5a + NOGGIN; (7) same as condition 5a.

Prior to experiments, reporter-line iBCs (region-specific and CSC protocol–derived) were sorted by NKX2-1 and TP63 reporter fluorescence plus NGFR staining (APC–anti-NGFR, Cat. 345108; BioLegend). Notably, for region-specific iBCs at D120 or later, sorting is often unnecessary because the majority are NGFR⁺. Non-reporter iBCs were sorted by NGFR and EpCAM (Alexa Fluor 488–EpCAM, Cat. 324210; BioLegend). All antibodies were used at 1:200 dilution. Sorted non-reporter iBCs can be expanded, maintained, or cryopreserved while remaining predominantly NGFR⁺; further sorting is not needed.

### ALI culture of iBCs or HBECs

Region-specific iBCs and CSC protocol-derived iBCs were seeded on 6.5 mm polyester membrane Transwell inserts with 0.4 μm pore size (Corning) coated with Matrigel diluted in DMEM/F12 (Gibco) at 200,000–300,000 cells per insert. Cells were seeded and maintained in their corresponding basal maintenance medium (1–5 as described above) for 7–12 days. Once Transwell membranes were covered by a dense confluent sheet of cells by visual inspection, the culture medium was switched to PneumaCult-ALI medium (STEMCELL Technologies) in both apical and basal chambers. The following day, the medium from top chamber was removed and the cells were further differentiated in ALI conditions for 2–3 weeks (for CSC protocol-derived iBCs) or 4 weeks (for region-specific iBCs) before analysis. HBECs were maintained and seeded onto Transwell inserts as described in previous paragraph, then switched to Vertex ALI or PneumaCult-ALI medium for 4 weeks before analysis.

### Flow cytometry analysis

Flow cytometry analysis was performed on a BD FACS LSR II, FACSMelody, or FACSAria II (BD Biosciences, San Jose, CA) at the UTHealth Flow Cytometry Service Center. Definitive endoderm induction was assessed by surface expression of CXCR4, c-KIT and EpCAM. Endodermal cells were dissociated into single cells with 0.05% trypsin/EDTA, then stained with APC-CXCR4 (Cat# 306510, BioLegend), PE-c-KIT (Cat# 313204, BioLegend) and Alexa Fluor 488-EpCAM (Cat# 324210, BioLegend) in PBS containing 0.1% BSA and 0.2 mM EDTA for 30**–**45 min at 4 °C. Lung, airway, and basal progenitors, as well as iBCs, were analyzed based on reporter and/or surface marker expression detected by fluorochrome-conjugated primary antibodies as described above. All antibodies were used at 1:200 dilution. DAPI or PI was used to exclude dead cells, depending on the instrument used. Data were analyzed with FlowJo (BD Biosciences).

### Tracheal xenografts

We used a tracheal xenograft model to evaluate the *in vivo* differentiation potential of proximal iBCs. Open-ended xenografts were assembled as previously described (Filali et al. 2002) with minor modifications: rat tracheas were decellularized by repeated freeze–thaw cycles and tubing was attached. Prior to seeding, xenografts were coated overnight at 4 °C with Matrigel diluted in DMEM/F12, then seeded with 1**–**2 × 10^6^ iBCs per trachea and allowed to attach overnight in culture medium at 37 °C. Two xenografts per mouse were implanted subcutaneously into the flanks of immunocompromised Nu/Nu mice, maintaining luminal air exposure via the open-ended tubing. After 4**–**5 weeks, grafts were harvested and analyzed by H&E staining and immunostaining.

### Immunofluorescence and Imaging

D15 lung progenitors grown in 2D were fixed in 4% paraformaldehyde (PFA) for 45 min at room temperature (RT). D30 airway/multipotent progenitors were recovered from 3D Matrigel droplets, dissociated to single cells, sorted for NKX2-1^GFP+^ cells, briefly plated on Matrigel-coated Transwell membranes for 24–48 h, then fixed in 4% PFA for 30 min at RT. OCT-embedded human fetal lung (HFL) tissue was cryo-sectioned at 6 µm on a Leica CM1950 cryostat and fixed in 4% PFA for 20 min. After fixation, D15 cells, D30 cells, and HFL sections were washed in PBS and permeabilized for 1 h, 30 min and 30 min, respectively, in PBS containing 0.25% Triton X-100 (MilliporeSigma) and 2% normal donkey serum (Jackson ImmunoResearch Labs, West Grove, PA). Samples were blocked in PBS with 2% donkey serum, 0.2% BSA, and 0.1% Tween-20 (blocking/staining buffer) for 1 h, incubated with primary antibodies in staining buffer overnight at 4 °C, then with appropriate secondary antibodies for 1 h at RT. Nuclei were counterstained with DAPI in PBS for 5 min. D15 cells were stored in PBS with ProLong Gold Antifade Mountant (Thermo Fisher) at 4 °C. D30 Transwell membranes were carefully excised, mounted in ProLong Gold Antifade Mountant with coverslips, and stored at 4 °C. HFL sections were mounted in ProLong Gold Antifade Mountant, coverslipped, and stored at 4 °C.

For whole-mount staining of ALI cultures, cells on Transwell inserts were fixed in 4% PFA for 60–80 min at RT, washed with PBS, permeabilized in PBS containing 0.5% Triton and 2% donkey serum for 3 h at RT, and blocked in blocking buffer for 1 h. Samples were incubated with primary antibodies in staining buffer with or without 0.1% Triton overnight at 4 °C, followed by appropriate secondary antibodies for 2 h at RT. Nuclei were counterstained with DAPI in PBS for 10 min. The Transwell membranes were then carefully excised, mounted in ProLong Gold Antifade Mountant on slides, and stored at 4 °C.

For paraffin embedding, ALI cultures and tracheal xenografts were fixed in 4% PFA overnight at 4 °C, washed in PBS, and processed by the UTHealth Microscopy Core. After graded ethanol dehydration, samples were cleared in xylene, infiltrated with paraffin, and embedded in wax. Sections (5–6 µm) were cut on an HM325 rotary microtome (Thermo Fisher). Two sections per sample were stained with hematoxylin and eosin. For immunofluorescence, sections underwent antigen retrieval with a citric acid–based unmasking solution (Vector Laboratories, Newark, CA), were permeabilized in PBS with 0.25% Triton and 2% donkey serum for 30–60 min, blocked for 1 h at RT, incubated with primary antibodies in staining buffer overnight at 4 °C, then with appropriate secondary antibodies for 1 h at RT. Nuclei were counterstained with DAPI in PBS for 5 min. Samples were mounted in ProLong Gold Antifade Mountant, coverslipped, and stored at 4 °C.

Samples were imaged on a Leica DMi8 automated inverted microscope (Leica Microsystems, Wetzlar, Germany) or a Nikon AX-R automated inverted confocal microscope (Nikon, Melville, NY). Tile scans were acquired using the tiling module with autofocus and auto-stitched in Leica Application Suite. Ionocytes were quantified manually by counting FOXI1- or BSND-positive cells in 40× transverse sections or 20× fields of whole-mount Transwell membranes. For each condition, ∼1,000 cells from random 40× transverse sections were analyzed, and frequencies were calculated relative to total cells (DAPI-stained nuclei). Ten random 20× fields per condition were quantified on whole-mount membranes, and total cells per field were recorded.

### Epithelial cell electrophysiological analysis

HBECs and proximal iBCs were differentiated in PneumaCult-ALI medium for 4 weeks prior to Ussing chamber analysis; CSC protocol–derived iBCs were differentiated for 2.5 weeks. Transepithelial ion currents of cell monolayers were assessed using EasyMount Ussing Chamber Systems operated under voltage-clamp mode, with tracings recorded and data analyzed using Acquire & Analyze software (Physiologic Instruments, Venice, FL). Briefly, Transwell inserts were mounted into the chambers and assayed under asymmetrical chloride conditions, with cell monolayers bathed on the apical side in low-chloride Ringer’s solution (1.2 mM NaCl, 140 mM sodium gluconate, 25 mM NaHCO₃, 3.33 mM KH₂PO₄, 0.83 mM K₂HPO₄, 1.2 mM CaCl₂, 1.2 mM MgCl₂, 10 mM glucose) and on the basolateral side in standard Ringer’s solution (120 mM NaCl, 25 mM NaHCO₃, 3.33 mM KH₂PO₄, 0.83 mM K₂HPO₄, 1.2 mM CaCl₂, 1.2 mM MgCl₂, 10 mM glucose).

Prior to measurements, a blank Transwell insert was mounted in the Ussing chamber with low-chloride Ringer’s solution on the apical side and standard Ringer’s solution on the basolateral side to compensate for electrode asymmetry and fluid resistance. The blank was then replaced with a cell-seeded insert, which was voltage-clamped and monitored for short-circuit current; transepithelial electrical resistance (TEER) was calculated using Ohm’s law at the time of baseline stabilization.

After baseline stabilization, amiloride (100 µM; MilliporeSigma) was added to both sides to inhibit epithelial sodium channel (ENaC) activity. Subsequently, forskolin (10 µM; MilliporeSigma) was applied to stimulate CFTR-mediated chloride current. At the end of recordings, CFTRinh-172 (10 µM; MilliporeSigma) was added apically to specifically inhibit CFTR, followed by apical UTP (100 µM) to assess epithelial integrity. The resulting changes in short-circuit current were calculated as ΔIsc and reported as mean ± SD.

### Single-cell RNA-sequencing and analysis

D45, 75 and D100 organoids were extracted, dissociated to single cells, live cells were sorted, and ∼10,000 cells per sample were prepared for scRNA-seq or sc-multiome. For ALI samples prepared at UTHealth, HBECs and D130 iBCs from Proximal 1, Proximal 2, Intermediate, and CSC conditions were differentiated in PneumaCult-ALI for 4 weeks (2.5 weeks for CSC), dissociated to single cells, and ∼10,000 live sorted cells per sample were processed for scRNA-seq. ALI samples processed at the CFF laboratory followed published methods (Carraro et al. 2021). scRNA-seq/sc-multiome for D45, 75, D100, and UTHealth ALI samples was performed at the MD Anderson Advanced Technology Genomics Core (ATGC) and the UTHealth Cancer Genomics Core. Single-cell suspensions (for scRNA-seq) or isolated nuclei (for sc-multiome) were processed using the 10x Genomics Chromium platform according to the manufacturer’s instructions. scRNA-seq libraries were generated using the Chromium Single Cell 3′ kit, and sc-multiome libraries using the Chromium Single Cell Multiome ATAC + Gene Expression kit, followed by sequencing on an Illumina NovaSeq 6000 or NovaSeq X Plus platform. For the sc-multiome dataset generated in this study, only the gene expression component was analyzed (Figure S2C-D). Data were processed with Cell Ranger, and downstream analyses were conducted in Seurat (R) with custom scripts. Ambient RNA was removed with SoupX; dead cells were excluded based on high mitochondrial RNA content and low/absent EPCAM; potential doublets and high-complexity outliers were removed based on the distribution of detected genes per cell. After QC, cells were classified using combinations of canonical airway lineage markers and visualized by UMAP. Replicates from the same condition (e.g., three Proximal 1 ALI and two Proximal 2 ALI samples) were integrated using Seurat’s unsupervised integration workflow. Dot plots, heatmaps, and feature plots of selected genes used log-normalized expression. Correlation analyses employed the top 4,500 highly variable genes per dataset. Seurat projects were kindly provided for three datasets (Carraro et al. 2021; Guo et al. 2023; Kadur Lakshminarasimha Murthy et al. 2022) and classifications for the remaining datasets (Basil et al. 2022; Okuda et al. 2020) were reproduced from published annotations. Raw data are deposited in GEO and will be available upon publication.

### Quantitative real-time PCR

Total RNA was extracted with the RNeasy Kit (Qiagen, Germantown, MD). RNA concentration was measured on a NanoDrop 2000 spectrophotometer (Thermo Fisher). For each sample, cDNA was synthesized from 1 µg total RNA using SuperScript IV First-Strand Synthesis with random hexamers (Invitrogen) in a 20-µl reverse-transcription reaction, according to manufacturer’s instruction. The resulting cDNA was used for multiple qPCR reactions, with each sample run in triplicate and each technical replicate containing cDNA corresponding to 10-20 ng input RNA. Quantitative real-time PCR was performed on an ABI 7900 thermocycler (Applied Biosystems) with Power SYBR Green PCR Master Mix. Cycling conditions were 50 °C for 2 min, 95 °C for 10 min, then 40 cycles of 95 °C for 15 s and 60 °C for 1 min, followed by melt-curve analysis. Relative expression was calculated by normalization to β-ACTIN and GAPDH, with reactions run in triplicate. Primer sets were previously published (Huang et al. 2014; Huang et al. 2015; Eenjes et al. 2021).

### Chromatin immunoprecipitation (ChIP)

Bulk cultures of hPSC-derived D6 anterior foregut endoderm (AFE) and D17 lung progenitors were crosslinked in 1% formaldehyde, quenched with 1 M glycine (pH 5) to a final concentration of 125 mM for 10 min at RT with gentle shaking, and processed for ChIP, library preparation, and analysis as previously described (Hassan and Chen 2024; Little et al. 2019). Rabbit anti-NKX2-1 (1 µg per reaction, ab133737, Abcam) was previously validated (Little et al. 2019) and used for experiments. Libraries were sequenced on an Illumina NextSeq 500 at MD Anderson ATGC. Reads were assessed with FastQC (v0.11.8), filtered and trimmed with Trimmomatic (v0.33), and aligned using Bowtie (parameters: -m1 -k1 -v1). SAM files were converted to BAM with SAMtools (v1.15) and filtered for unmapped/low-quality reads using Picard MarkDuplicates (v2.9.0) and SAMtools (settings: “-b -h -F 4 -F 1024 -F 2048 -q 30”). Peaks were called with MACS2 (v2.4.1) and filtered against the mm10 blacklist. Peaks were assigned to the nearest gene using ChIPseeker (v1.3).

### Quantification and Statistical Analysis

We include at least three biological replicates per experiment and four mice per condition (eight xenografts, 60–70% success rate), meeting power requirements (α = 0.05, 1 − β = 0.8; LaMorte’s Biostatistics Tool). Statistical significance was evaluated using parametric tests (unpaired/paired two-tailed Student’s t-test or ANOVA). Data are presented as mean ± SD; p < 0.05 was considered significant.

## ACKNOWLEDGMENTS

We are grateful to Dr. Scott Randell and the CF Center Tissue Procurement and Cell Culture Core at the University of North Carolina for generously providing HBECs. We thank Drs. Yan Xu and Minzhe Guo at Cincinnati Children’s Hospital Medical Center for kindly providing the Seurat object for the LungMAP CellRef meta-dataset. We thank Dr. Ville Meretoja at the University of Texas Health Science Center at Houston (UTHealth) Flow Cytometry Core for assistance with sample sorting, and Dr. Zhengmei Mao of the UTHealth Microscopy Core Facility for histological sample preparation. We acknowledge UTHealth Cancer Genomics Core, supported by the Cancer Prevention and Research Institute of Texas (CPRIT) grant RP240610, and the MD Anderson Cancer Center Advanced Technology Genomics Core Facility, supported by NIH grants CA016672 and 1S10OD024977-01. This work was supported by NIH grants R01AR077045 and R21AR079075 to N.N.; NIH/NIA grant R01AG082132 to W.L.; NIH grant R01HL159712 and Department of Defense (DoD) grant W81XWH2110081 to Y.-W.C.; Chan Zuckerberg Initiative Seed Network (2019-002440) and Pediatric Network (2021-237566), an advised fund of the Silicon Valley Community Foundation, NIH/NHLBI grants R01HL119215 and R01HL166139, and CFF grant 008697225 to J.R.S.; NIH R01CA246130 and DoD RCRP HT9425-24-1-0957 to D.-F.L.; NIH grant R01HL139876 to B.R.D. and E.J.S.; NIH grant R01HL153511 to J.C.; Cystic Fibrosis Foundation (CFF) grant DAVIS20XX2 to B.R.D.; and UT Rising STARS Award and CFF grants HUANG22G0 and HUANG24P0 to S.X.H.

## AUTHOR CONTRIBUTIONS

S.X.H. conceived the study. S.S., J.C., B.R.D., and S.X.H. developed the differentiation strategy, designed experiments, analyzed data and wrote the manuscript. S.S., N.V.AS, S.D., B.C.D, S.L.W, C.B. and N.F-C. performed differentiation experiments. S.S. and B.R.D. designed the targeting strategy and generated the TP63-tdTomato hiPSC. C.M.L designed the targeting strategy and generated the SOX2-mRuby3 hPSC. J.L. and J.E.M provided the TP63-mCherry and CF G551D hiPSCs and conducted the CFTR protein characterization experiments. D.M.H., M.-F.H, B.T. and A.D.C. performed bioinformatic analysis of scRNA-seq, sc-multiome and ChIP-seq data. P.P.H, T.F., and J.R.S. provided human fetal lung scRNA-seq datasets. A.R., S.S.P, and E.J.S. performed Ussing chamber analysis. M.G.G., J.K., N.N., W.L., D.-F.L., and K.S.R. provided critical input. M.R-M. and Y.-W.C. prepared and provided human fetal lung tissues. J.M.A performed tracheal xenograft experiments.

