## Supplementary Figures and Legends for "Generation of Region-specific Airway Basal Stem Cells from Human Pluripotent Stem Cells via Regulation of NOGGIN-BMP Axis"

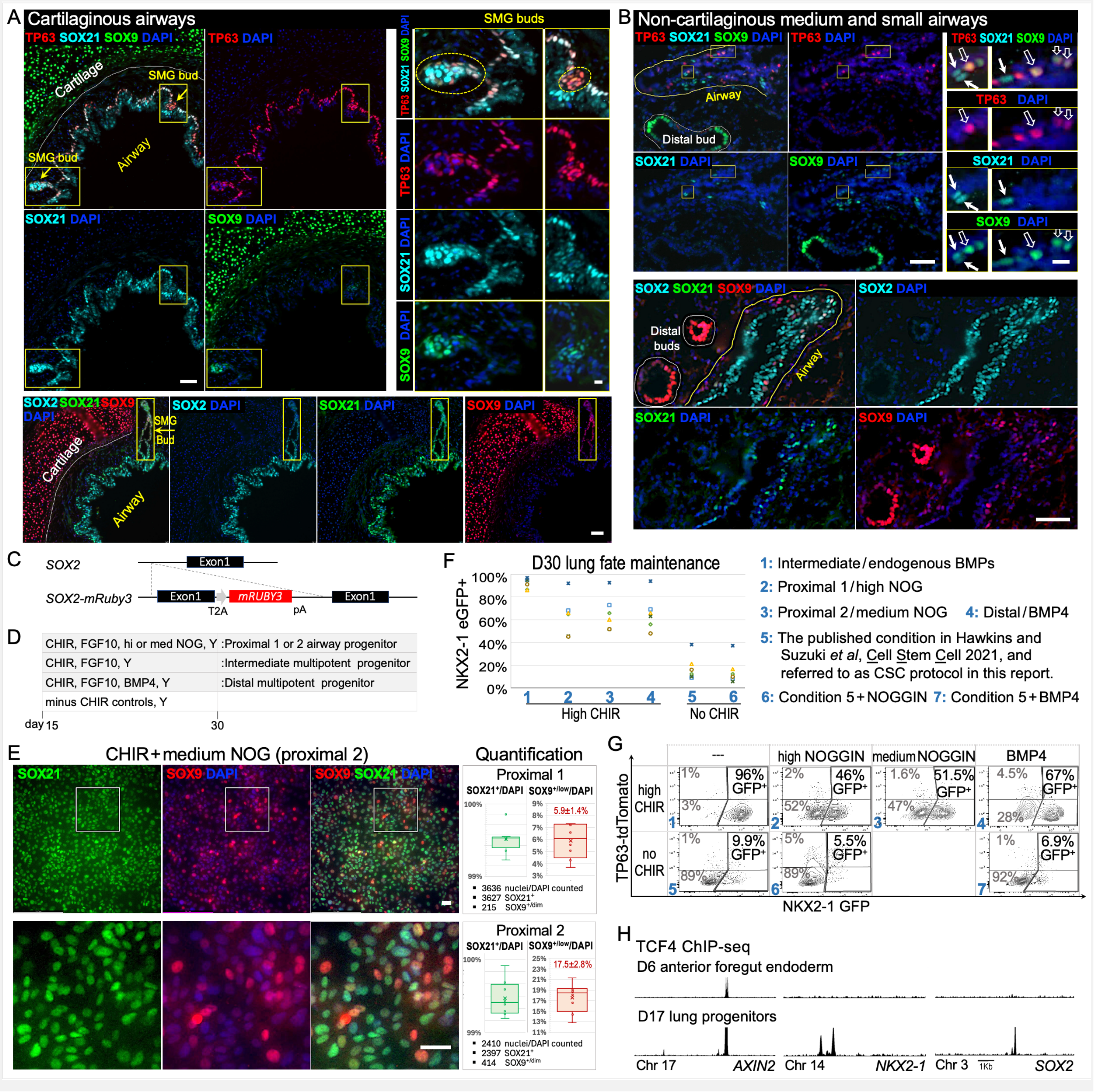


**Supplementary Figure 1**. **Characterization of SOX2, SOX21, and SOX9 expression in human fetal airways and NKX2-1 lung fate maintenance in D30 hPSC-derived lung progenitors**. Human fetal lungs from gestational weeks 16–18 were analyzed, and representative results from a GW16.5 lung are shown in A and B. (**A**) Cartilaginous airways contain a homogenous population of $SOX2^{+}SOX21^{+}SOX9^{-}$ cells. $TP63^{+}$ basal progenitors are abundant along the basal lamina (top panels). The only $SOX9^{+}$ small cell clusters are budding submucosal glands (top panels, insets). **(B)** In non-cartilaginous small airways from GW16.5 lungs, $TP63^{+}$ cells are rare. These regions show uniform SOX2, with sparser SOX21 and SOX9 expression. $SOX21^{+}SOX9^{+}$ (closed arrows) and $TP63^{+}SOX9^{+}$ (open arrows) cells are observed (top panels, insets). (**C**) Schematic illustration of SOX2-mRuby3 reporter construct. (**D**) Schematic of the days 15–30 culture protocol (stage 2). CHIR:

a pharmacological agonist of WNT signaling. (**E**) *Left*, SOX21 and SOX9 immunofluorescence in NKX2-1$^{GFP+}$ cells from the CHIR + medium NOG condition, cultured according to D. Cells were isolated from 3D culture, NKX2-1$^{GFP+}$ cells were sorted, and briefly plated in 2D for 24–48 hours prior to staining. The bottom panel shows an enlarged inset. Scale bar, 50μm. *n* = 3 for each hPSC line; representative images from the BU3 NGPT hiPSC line. *Right*, quantification of SOX21$^{+}$ and SOX9$^{+/low}$ cells under proximal 1 (Figure 1G, *left)* and proximal 2 (*left* panel) conditions. (**F**) NKX2-1 expression was assessed by GFP reporter protein levels using FACS in the conditions cultured according to D, with and without WNT activation through CHIR. NKX2-1 lung fate at D30 as quantified as the percentage of GFP$^{+}$ cells from two biological replicates of each of the three reporter lines. *n* = 2-4 for each hPSC reporter line. Paired *t* test, condition 1 versus 5, P<0.001; condition 2 versus 5, P<0.005. (**G**) A representative example from BU3 NGPT of experiments presented in F. Results from conditions 5-7 demonstrate that WNT activation through CHIR is required for lung fate maintenance independent of NOGGIN-BMP signaling, and that premature WNT withdrawal leads to undesired NKX2-1 lung fate loss. (**H**) TCF4 binds to 3′ sites of *NKX2-1* and *SOX2* in hPSC-derived D17 lung progenitors but not in their foregut precursors, whereas TCF4 binding is similar near a positive control gene *AXIN2*.

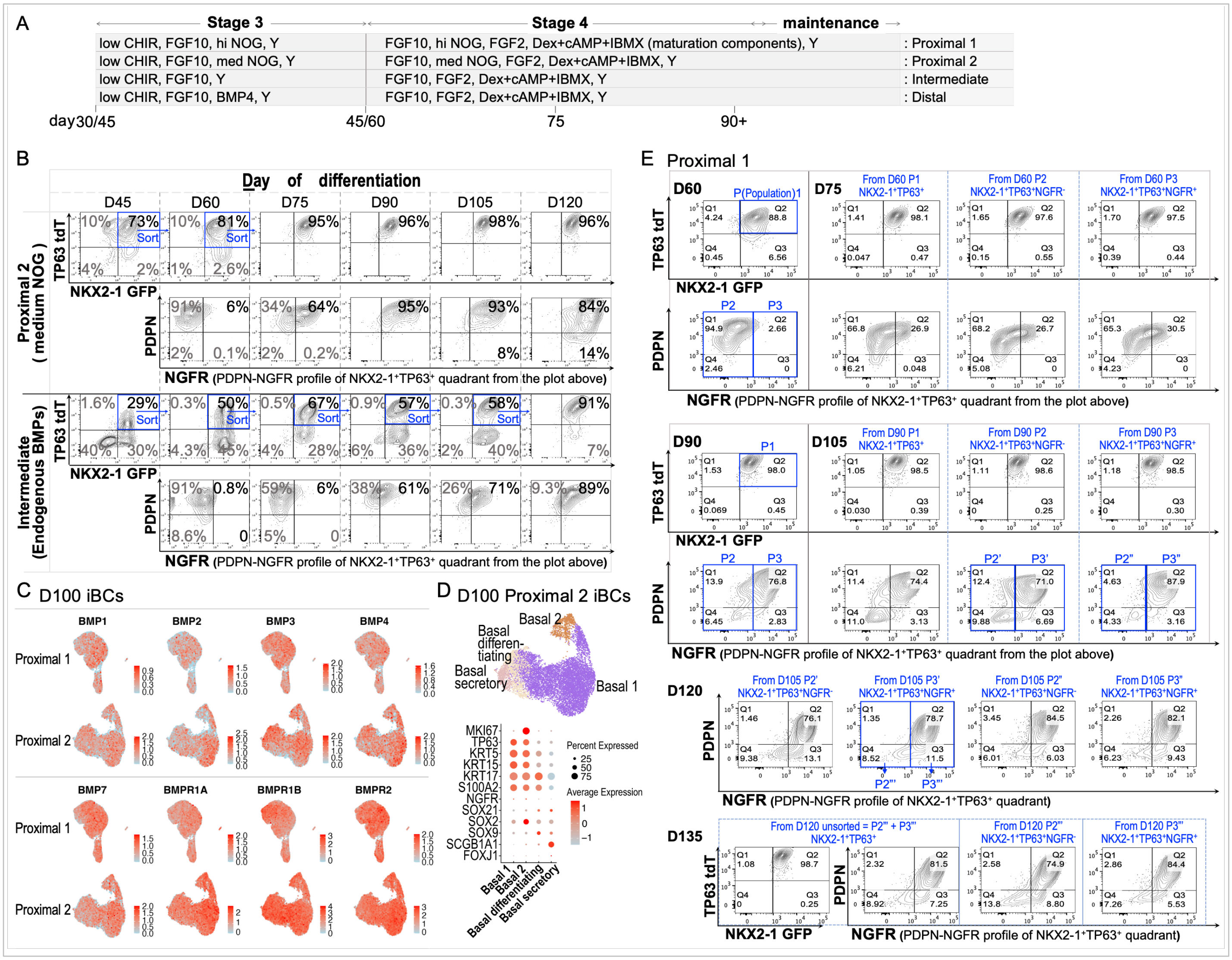


**Supplementary Figure 2. Generating of regio- specific iBCs (proximal 2 and intermediate) and characterization of BMP signaling and NGFR expression in proximal iBCs.** (**A**) Schematic of the stages 3–4 culture protocol from days 30/45 to 90+ (D30–90+ for proximal and intermediate conditions; and D45–105+ for the distal condition). Dex, dexamethasone; cAMP, cyclic AMP; IBMX, isobutylmethylxanthine. (**B**) Basal progenitor specification, maintenance, and maturation in proximal 2 and intermediate conditions. Basal progenitor specification efficiency was measured as the percentage of NKX2-1$^{GFP+}$ TP63$^{tdT+}$ cells at D45; basal maintenance during D60-120 was assessed as the percentage of cells that remain GFP$^{+}$tdT$^{+}$ at each time point; basal maturation was measured by NGFR expression in GFP$^{+}$tdT$^{+}$ cells. *n* = 3 for each hPSC line; a representative experiment from BU3 NGPT showing proximal 2 and intermediate conditions. Notably, some proximal NGFR$^{+}$ iBCs lost PDPN at D105-120. (**C**) Feature plots of BMP pathway signatures and receptor expression in D100 proximal BU3 NGPT cells by single-cell analysis; corresponding UMAPs are shown in Figures 2F and S2D. (**D**) Molecular signatures of basal (TP63, KRT5, KRT15, KRT17, S100A2, NGFR), proliferative (MKI67), proximal-distal patterning (SOX2, SOX21, SOX9), secretory (SCGB1A1), and ciliated (FOXJ1) markers in D100 proximal 2 cells from BU3 NGPT assessed by scRNA-multiome. UMAPs (*top*) display the identified clusters, and corresponding dot plots (*bottom*) show expression of the indicated markers across clusters. (**E**) Tracking of NGFR$^{+}$ iBCs and NGFR$^{-}$ basal progenitors under the proximal 1 condition by serial NGFR-based FACS. At D60 NGFR$^{+}$ and NGFR$^{-}$ fractions were sorted and cultured separately in 3D. Every 15 days, each culture was analyzed, NGFR$^{+}$ and NGFR$^{-}$ sub-populations were sorted and cultured separately as shown. This cycle was repeated through D135 (five successive NGFR sorts), and results from five rounds are shown.

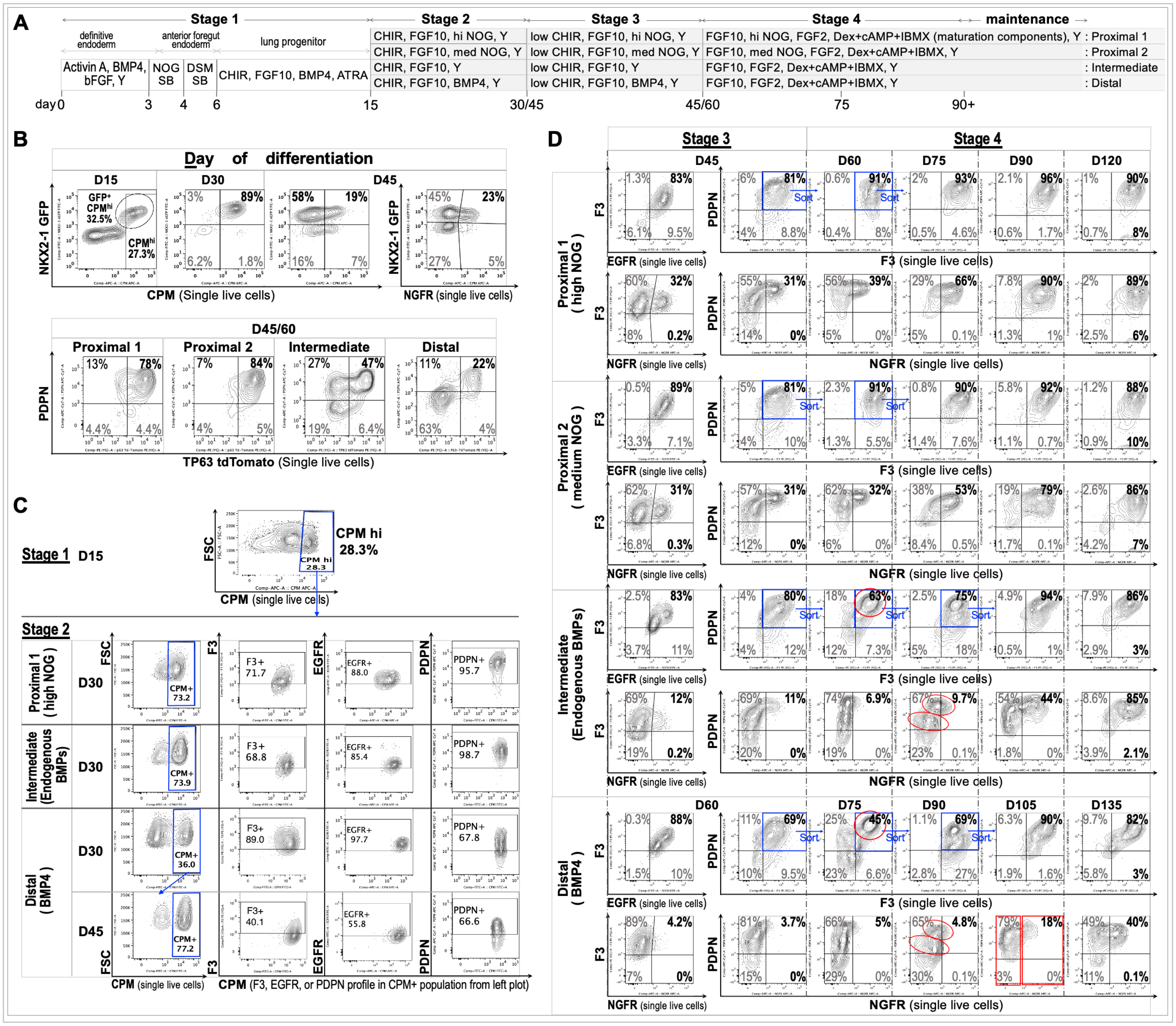


**Supplementary Figure 3. Sorting strategy for generating region-specific iBCs in non-reporter hPSCs.** (**A**) Schematic of days 0–90+ culture protocol. (**B**) Validation of surface markers for sorting using reporter lines. *Top*: under full WNT activation through CHIR, $CPM^{hi}$ and $CPM^{+}$ populations closely correlate with NKX2-1$^{GFP+}$ cells at D15 and D30, respectively. A representative D30 result from the intermediate condition is shown. After CHIR reduction, CPM expression is no longer specific to $GFP^{+}$ cells, whereas $NGFR^{+}$ cells arise exclusively from the $GFP^{+}$ fraction; representative D45 plots from the proximal 2 condition are shown. *Bottom*: PDPN enriches TP63$^{tdT+}$ populations across all conditions at the end of stage 3 basal specification (D45 for proximal and intermediate conditions; and D60 for the distal condition). (**C**) Purification of $CPM^{hi}$ lung progenitors at D15 for specification into D30 proximal airway progenitors and intermediate/distal multipotent progenitors, together with F3, EGFR, and PDPN profiles at D30. *n* = 3 for each non-reporter hPSC line; a representative experiment from RUES2 is shown. (**D**) Basal progenitor specification (stage 3), maintenance, and maturation (stage 4) under the indicated conditions. F3-EGFR and F3-NGFR profiles are shown for single live cells at D45 for proximal and intermediate conditions and at D60 for the distal condition (stage 3), whereas PDPN-F3 and PDPN-NGFR profiles are shown throughout stages 3–4. At stage 3, $NGFR^{+}$

cells were detected predominantly within the F3$^+$ and PDPN$^{hi}$ fractions of single live cells (bottom panels of each condition), supporting the use of PDPN$^+$F3$^+$ sorting to enrich basal progenitors at D45–60 under proximal and intermediate conditions and at D60–75 under the distal condition. Under intermediate and distal conditions, additional PDPN$^+$F3$^+$ sorting was performed at D75 and D90, respectively. Under these conditions, a more stringent alternative strategy, indicated by red oval gates, involved isolating PDPN$^{hi}$F3$^{hi}$ populations at D60 and D75, respectively, followed 15 days later by separation of PDPN$^{hi}$ and PDPN$^+$ populations (at D75 and D90, respectively) for further culture. At D75+ under the intermediate condition and D90+ under the distal condition, red rectangular gates indicate separation of NGFR$^+$ and NGFR$^-$ populations for culture, which promoted NGFR expression in the NGFR$^-$ fraction, as shown in Figure S2E. *n* = 3 for each non-reporter hPSC line; a representative experiment from RUES2 is shown.

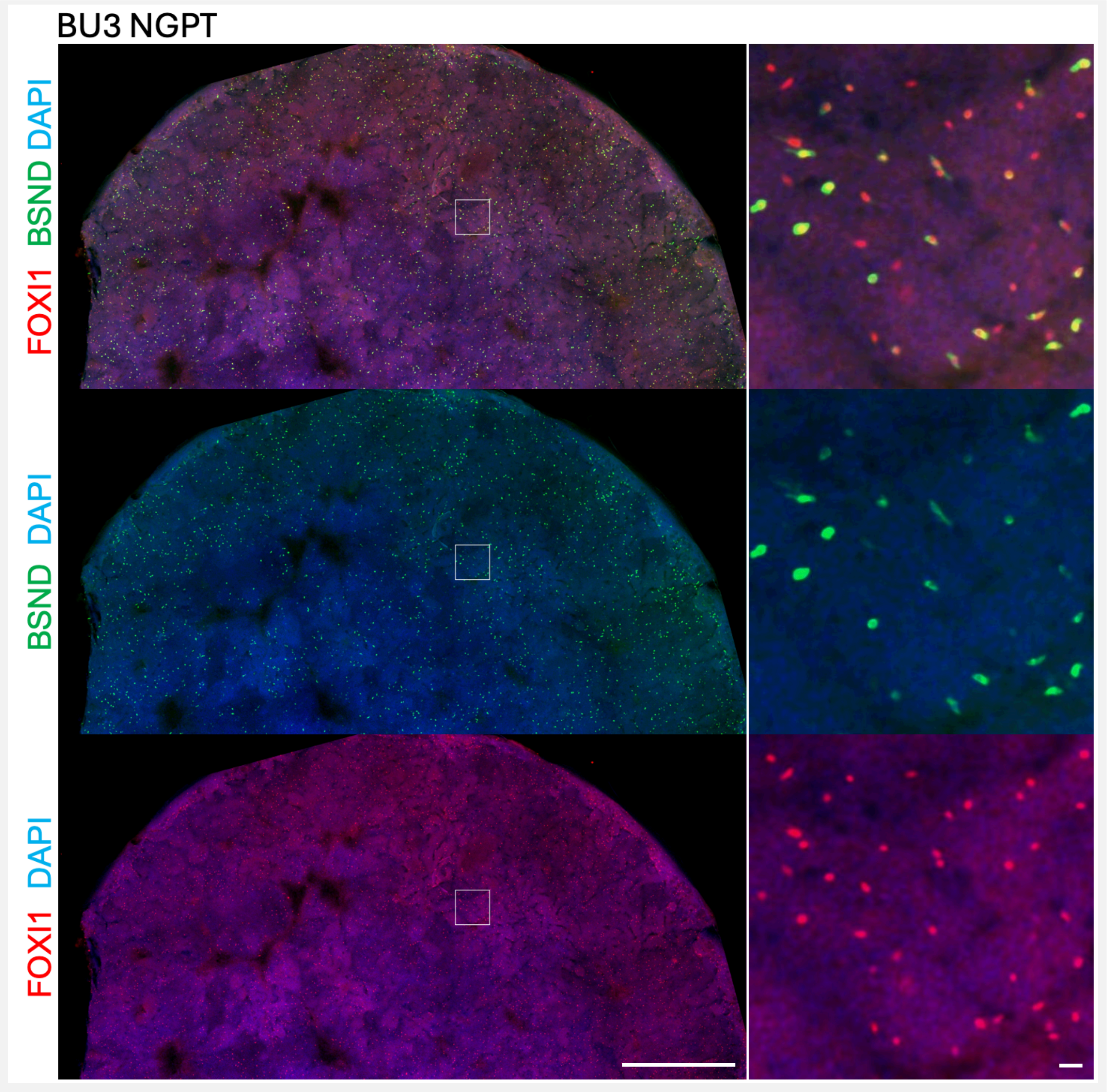


**Supplementary Figure 4. Ionocytes in proximal 1 iBC-ALI cultures derived from BU3 NGPT hiPSCs.** D90–130 proximal 1 iBCs were differentiated in ALI medium for 4 weeks and stained for FOXI1 and BSND. Representative images from $n$ = 3 experiments are shown. *Left panel*, tiled scan of half a Transwell; scale bar, 1mm. *Right panel*, enlarged view of the boxed inset; scale bar, 20μm.

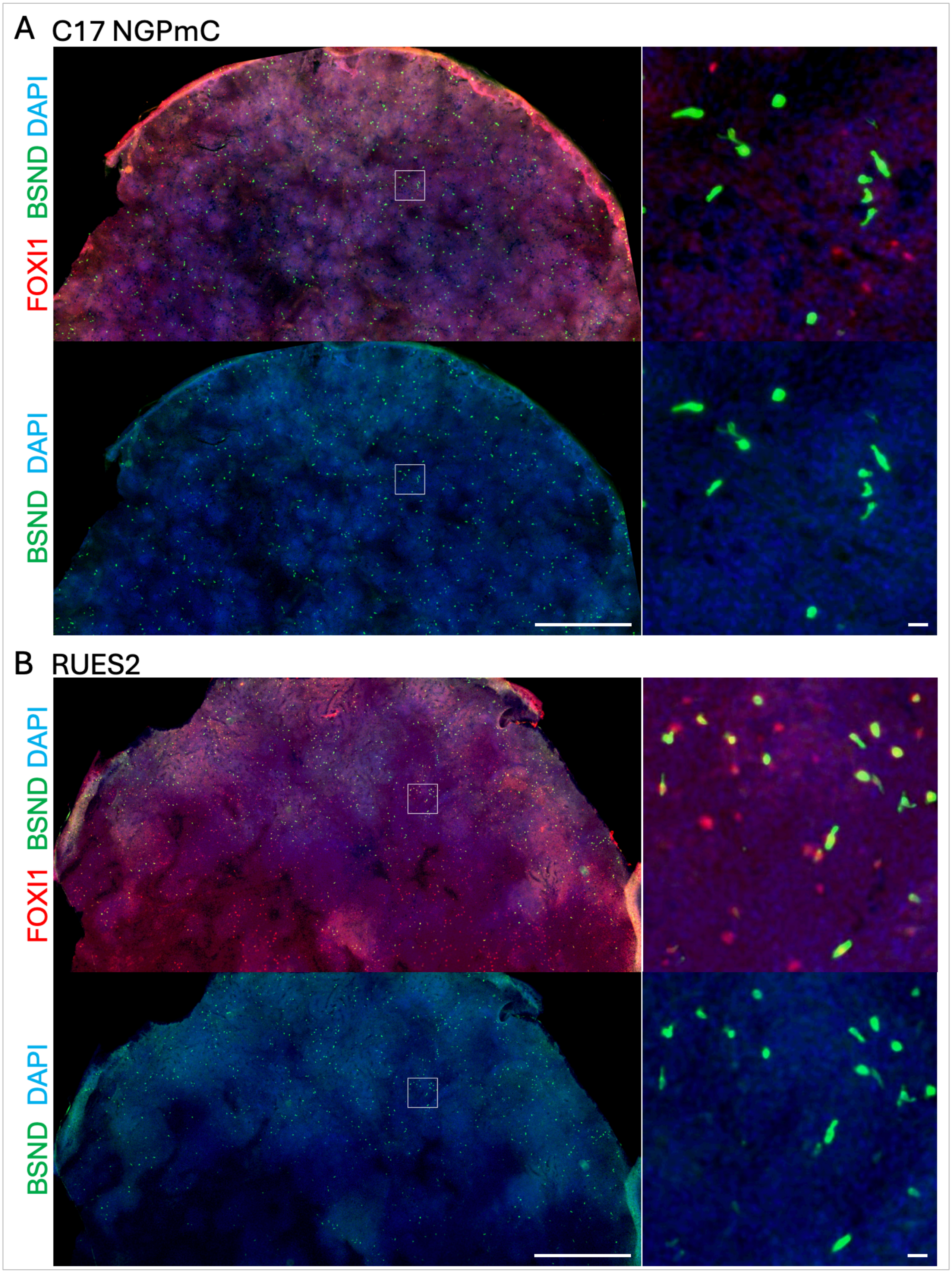
A
C17 NGPmC
FOXI1 BSND DAPI
BSND DAPI
B
RUES2
FOXI1 BSND DAPI
BSND DAPI

**Supplementary Figure 5. Ionocytes in proximal 1 iBC-ALI cultures derived from (A) C17 NGPmC hiPSCs and (B) RUES2 hESCs.** D90–130 proximal 1 iBCs were differentiated in ALI medium for 4 weeks and stained for FOXI1 and BSND. Representative images from $n$ = 3 experiments for each line are shown. *Left panels*, tiled scan of half a Transwell; scale bar, 1mm. *Right panels*, enlarged view of the boxed insets; scale bar, 20μm.

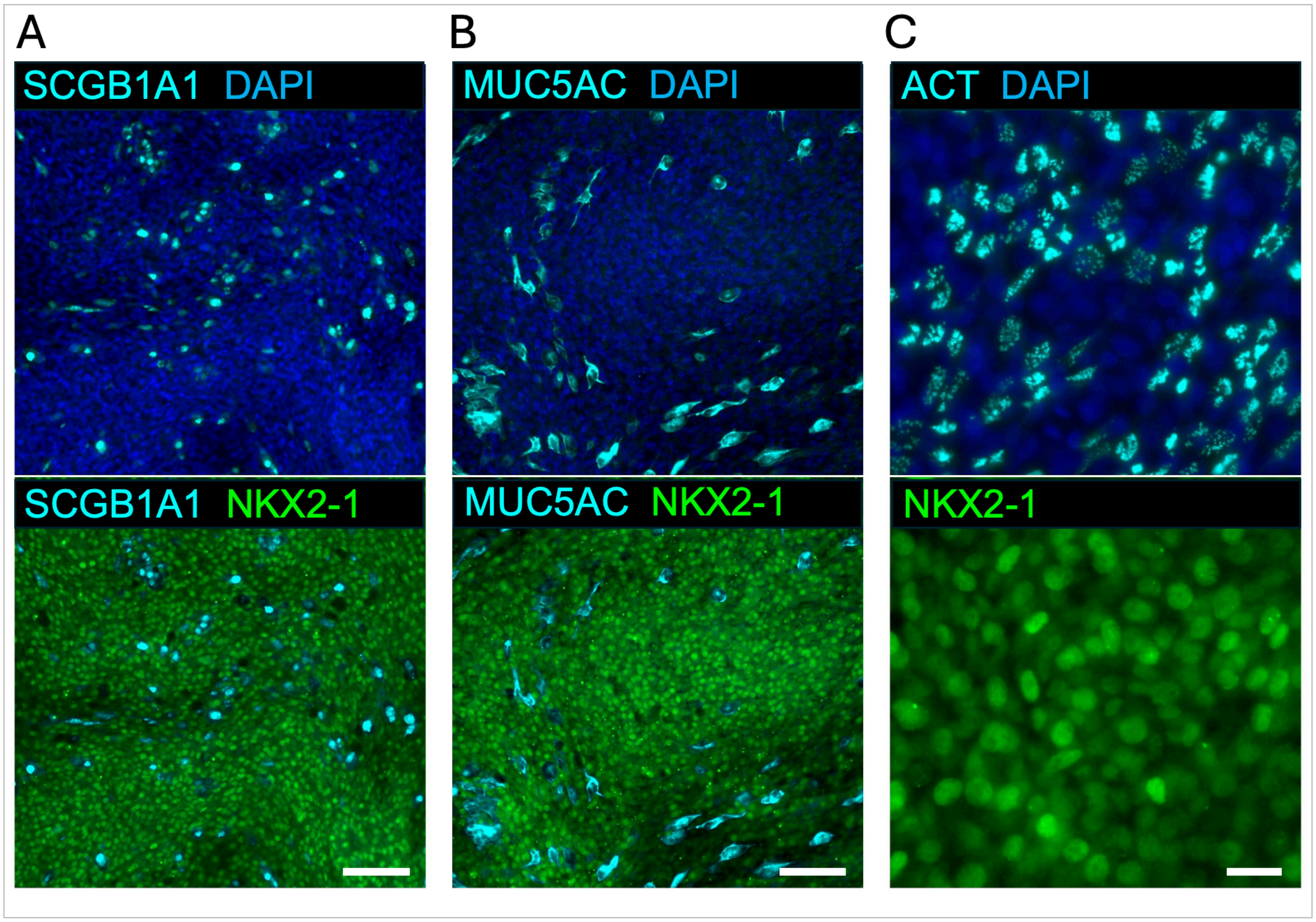


**Supplementary Figure 6. Secretory and ciliated cells in proximal iBC-derived ALI cultures.** D90–130 proximal iBCs were differentiated in ALI medium for 4 weeks and stained for (**A**) SCGB1A1/NKX2-1, (**B**) MUC5AC/NKX2-1, and (**C**) ACT/NKX2-1. $n$ = 3 for each hPSC line; representative images from BU3 NGPT hiPSC-derived proximal 1 iBC cultures are shown. Scale bars, 100μm in A and B, and 20μm in C.

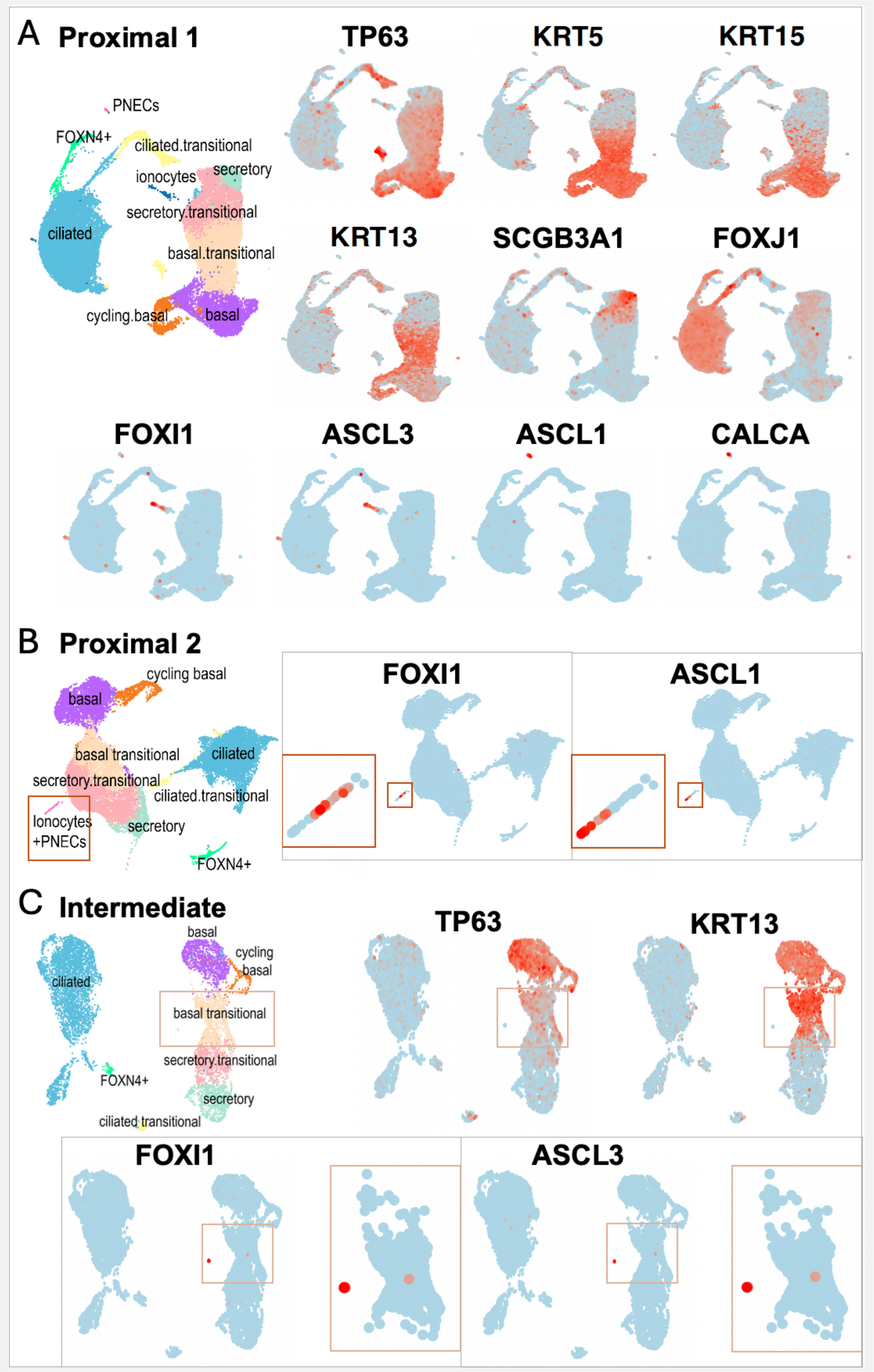


**Supplementary Figure 7. Feature-plot molecular signatures of region-specific iBC samples profiled by 10x Genomics scRNA-seq, as described in Figure 4. (A)** In proximal 1 samples, each lineage/cluster is clearly separated on the UMAP and distinguished by its key DEG(s), as shown in the feature plots. **(B)** In proximal 2 samples, ionocytes and PNECs cluster together on the UMAP; *FOXI1* and *ASCL1* feature plots with enlarged insets (lower left) show that the two lineages are adjacent but segregated. **(C)** Under the intermediate condition, a small subset of ionocytes overlaps spatially with basal transitional cells (*TP63*$^{low}$*KRT13*$^{hi}$); *FOXI1* and *ASCL1* feature plots with enlarged insets mark the few positive cells.

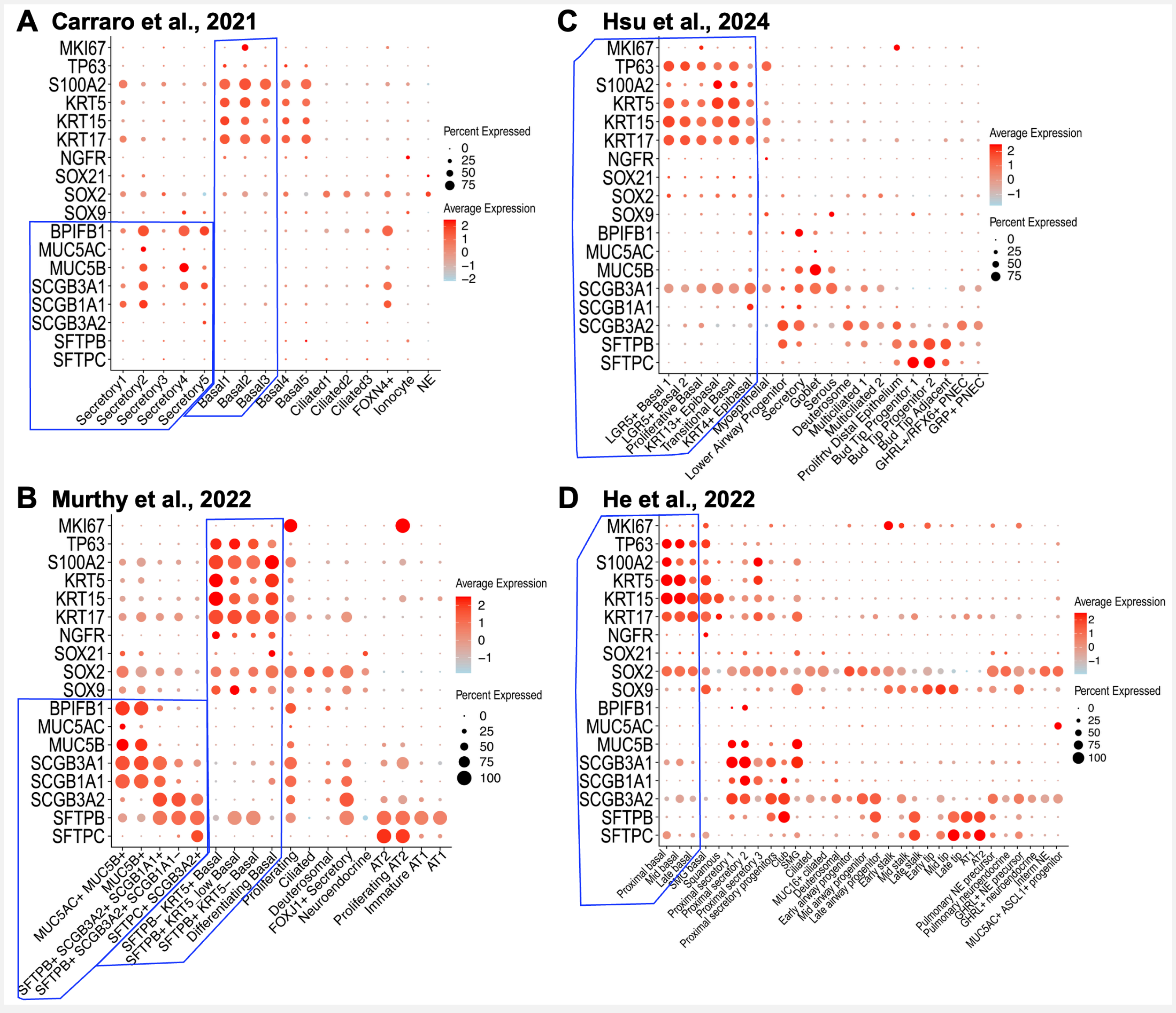


**Supplementary Figure 8. Dot plots of basal and secretory molecular signatures in adult and fetal airway/lung tissue datasets.** Reanalysis of **(A)** GSE150674 (Carraro et al., 2021, Figure 1C) and **(B)** GSE178360 (Kadur Lakshminarasimha Murthy et al., 2022, Figure 1C); blue rectangles indicate secretory and basal populations as defined by the authors. *SOX9* expression is stronger in distal airway basal populations than in proximal airway basal cells (B versus A). Reanalysis of **(C)** Hsu *et al*. (2024, Figure 2A; data shared by the authors through collaboration) and **(D)** He *et al*. (2022, Figure 2A; data available through the fetal lung cell atlas portal); blue rectangles indicate basal subpopulations as defined by the authors.
